# Differential modulation of miR-122 transcription by TGFβ1/BMP6: implications for nonresolving inflammation and hepatocarcinogenesis

**DOI:** 10.1101/2023.06.30.547174

**Authors:** Martha Paluschinski, Claus Kordes, Mihael Vucur, Veronika Buettner, Christoph Roderburg, Haifeng C Xu, Prashant Shinte, Philipp A Lang, Tom Luedde, Mirco Castoldi

## Abstract

Chronic inflammation is widely recognized as a significant factor that promotes and worsens the development of malignancies, including hepatocellular carcinoma. This study aimed to explore the potential role of microRNAs in inflammation-associated nonresolving hepatocarcinogenesis. By conducting a comprehensive analysis of altered microRNAs in animal models with liver cancer of various etiologies, we identified miR-122 as the most significantly downregulated microRNA in the liver of animals with inflammation-associated liver cancer. Although previous research has indicated the importance of miR-122 in maintaining hepatocyte function, its specific role as either the trigger or the consequence of underlying diseases remains unclear. Through extensive analysis of animals and *in vitro* models, we have successfully demonstrated that *MIR122* transcription is differentially regulated by the immunoregulatory cytokines by the transforming growth factor-beta 1 (TGFβ1) and the bone morphogenetic protein-

6 (BMP6). Furthermore, we presented convincing evidence directly linking reduced *MIR122* transcription to inflammation and in chronic liver diseases. The results of this study strongly suggest that prolonged activation of signaling pathways, leading to disruption of cytokine-mediated regulation of *MIR122*, may significantly contribute to the onset and exacerbation of chronic liver disease.

## Introduction

Persistent inflammation has emerged as a critical factor in the development and progression of malignancy [1]. In the context of the liver, unresolved inflammation can lead to a cascade of events, including hepatic injury, fibrosis, cirrhosis, and ultimately the onset of hepatocellular carcinoma (HCC; [2]). Regardless of the underlying causes, a common feature in HCC initiation is the perpetuation of a wound-healing response triggered by parenchymal cell death and subsequent inflammatory cascades. Therefore, gaining insights into the signaling pathways involved in the transition from acute to chronic liver injury and from dysplasia to HCC is crucial for the identification of potential predictive biomarkers and therapeutic targets in patients with chronic liver diseases.

microRNAs (miRNAs) are small RNA molecules that play key roles in regulating gene expression and controlling a wide range of cellular processes [3]. They are known to respond to cellular signals and pathological conditions by modulating gene expression. In this study, we aimed to identify miRNAs associated with non-resolving inflammation by conducting a high-throughput analysis of miRNA expression in HCC models characterized by an inflammatory milieu. Our results revealed significant downregulation of miR-122 in tumor tissues isolated from the livers of mice exhibiting a uncontrolled inflammatory state. miR-122 is known to play a central role in the regulation of diverse aspects of hepatic function, while its expression is often dysregulated in liver diseases, including fibrosis [4], cirrhosis [5], viral hepatitis [6,7], and HCC [8]. miR-122 knockout mice exhibit a hepatic phenotype characterized by liver inflammation, fibrosis, and a high incidence of liver cancer [9]. Notably, we have previously demonstrated that miR-122 controls iron homeostasis, and its inhibition in mice leads to systemic iron deficiency [10]. Specifically, miR-122 regulates hepcidin (Hamp) by modulating the translation of hemochromatosis (Hfe) [11] and hemojuvelin (Hjv) [12], which are crucial upstream activators of Hamp transcription. Furthermore, we have previously observed a significant reduction in miR-122 expression in the livers of Hfe knockout animals and in liver biopsies from patients with hereditary hemochromatosis (HH), suggesting the existence of an uncharacterized regulatory loop. Notably, Hamp, in addition to its role in iron metabolism, is known to play a critical role in linking the immune response to infection and inflammation [13]. In this study, we provide novel evidence indicating that the downregulation of miR-122 is a physiological response triggered by inflammation. Our findings suggest that the decrease in miR-122 expression may be necessary to redirect cellular pathways away from “housekeeping” functions such as iron homeostasis and contribute to the activation of the inflammatory response, hence shedding new light on the intricate relationship between miR-122, inflammation and chronic liver diseases.

## Results

### Integrative analysis of miRNAs expression in HCC models

In a previous study, we conducted an analysis of miRNA expression in the livers of mouse models with liver tumors of various etiologies [14]. These animal models included two cancer models characterized by an inflammatory environment, namely the transgenic over-expression of Lymphotoxin (AlbLTα/β) model and the c-myc transgenic mice (Tet-O-Myc mice) model, both resulting in inflammation-driven liver cancer. Additionally, we examined a non-inflammatory model where chemical hepatocarcinogenesis was induced using Diethylnitrosamine (DEN). Building upon this earlier work, our current research aimed to identify miRNAs that are globally associated with cancer and responsive to inflammation by analyzing changes in miRNA expression in distinct animal models of inflammation-driven liver cancer, specifically the AlbLTα/β and Tet-O-Myc mice. To achieve this, we obtained the raw data from our original study, which were downloaded from the Genome Omnibus repository (GEO, [15]; accession number: GSE102417) and reanalyzed the miRNA expression using the approach described in the materials and methods section. Our findings demonstrate a significant downregulation of miR-122 in the mouse models of liver cancer characterized by an inflammatory milieu (i.e., AlbLTα/β and Tet-O-Myc mice), while no significant change was observed in the livers of DEN-treated mice (**Figure 1A-C**). To independently evaluate these findings, we quantified miR-122 expression in the livers of transgenic mice lacking TNF receptor-associated factor 2 (*Traf2*) and Caspase-8 (*Casp8*) in liver parenchymal cells (LPC; LPC^Δ*Traf2*^, *^Casp8^*). These LPC^Δ*Traf2*^, *^Casp8^* mice were observed to develop spontaneous, immunogenic-driven liver tumors at 52 weeks of age (**Figure 1D-E**; [16]. Interestingly, a notable reduction in miR-122 expression was observed specifically in the liver of 6-week-old LPC^Δ*Traf2*^, *^Casp8^* mice, whereas no significant changes were observed in the livers of older animals (**Figure 1F**). Importantly, the levels of miR-122 were found unchanged in the liver of LPC^Δ*Traf2*^, *^Casp8^* –Ripk3^-/-^ and LPC^Δ*Traf2*^, *^Casp8, Ikbkb^* triple knockout mice, which were previously shown to rescue the LPC^Δ*Traf2*^, *^Casp8^* phenotype in both 6 and 52 weeks old mice (**Supplementary Figure S1A;** [16]). Although the true significance of this finding is so far unclear, it is plausible to speculate that the downregulation of this miRNA in the liver of LPC^Δ*Traf2*^, *^Casp8^* mice may contribute to establishment of a pro-inflammatory niche in the liver of these animals.

**Fig. 1.**
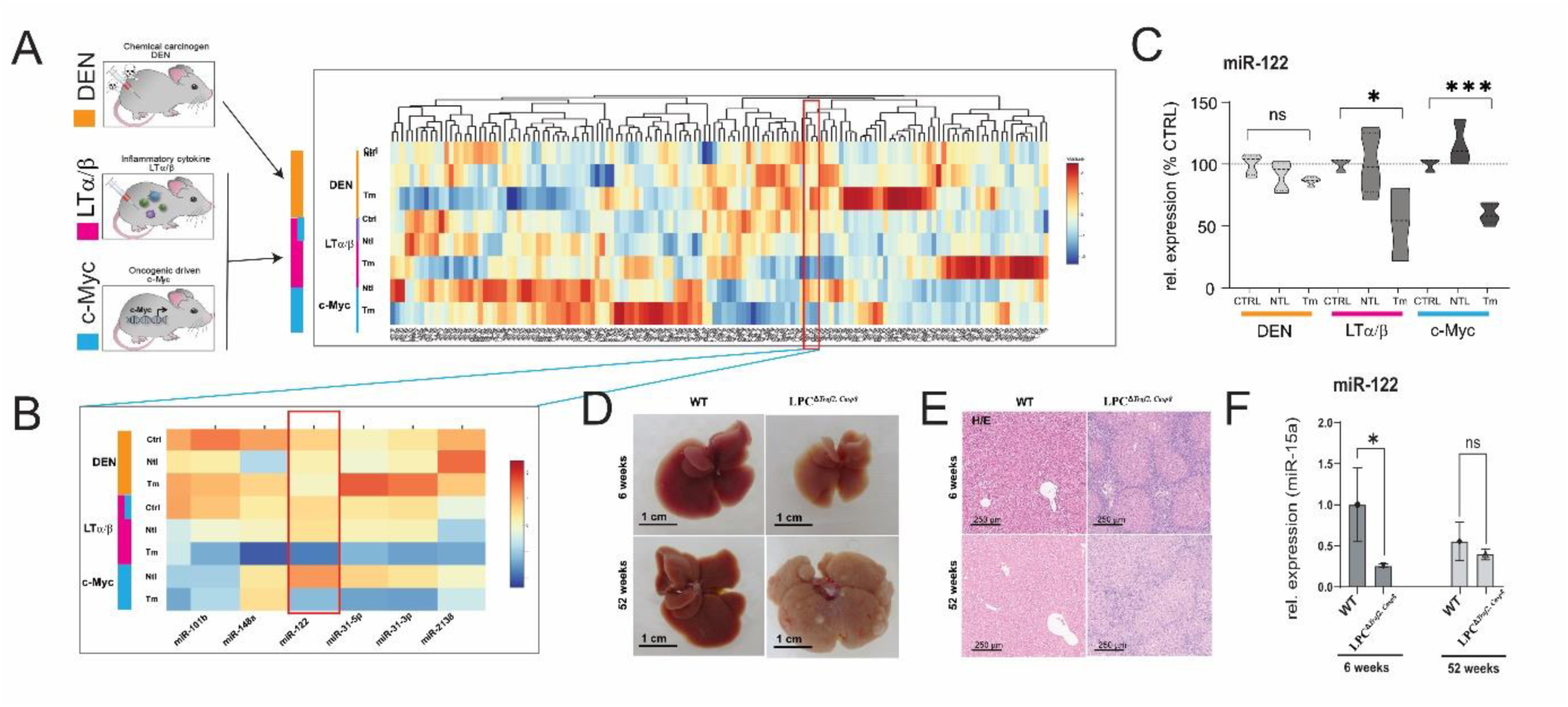
miR-122 is downregulated in inflammatory hepatocellular carcinoma. miR-122 is significantly downregulated in the tumor tissue isolated from the livers of lymphotoxin (LT, Purple) c-Myc (Blue) over-expressing animals but not in the tissue isolated from (DEN, Orange) animals. (**A**) Heat map depicting unsupervised hierarchical clustering of HCC models with respect to controls. The color scale illustrates the fold change of miRNAs across the samples. (**B**) Visualization of miRNAs downregulated in the tumor tissue of LT and c-myc models but not in the DEN model. (**C**) miR-122 expression as box plots (n = 5). (**D**) Representative image on the livers of wild type (ET) and LPC^Δ*Traf2*^, *^Casp8^* mice at 6 and 52 weeks. (**E**) Representative HE staining of the liver of control (WT) and LPC^Δ*Traf2*^, *^Casp8^* mice at 6 and 52 weeks of age. (**F**) Expression of miR-122 in the liver of WT and LPC^Δ*Traf2*^, *^Casp8^*mice at 6 and 52 weeks (normalized to miR-15a, n = 4). Results are represented as mean ± SD, significant differences were evaluated by using One-way ANOVA or t-test as appropriate (ns = Not significant, *, p ≤ 0.05; **, p ≤ 0.001; ***, p ≤ 0.0001).

### miR-122 expression is dysregulated in the liver of Hjv and Tmprss6 Ko mice

To investigate miR-122’s role in non-resolving acute inflammation, we analyzed the mechanisms behind its downregulation. We have previously shown that miR-122 was downregulated in the livers of Hfe deficient mice [10] suggesting that signaling pathways downstream to Hfe are required to maintain transcription of *MIR122* gene. Two signaling pathways are known to act downstream to Hfe: i) the extracellular signal-regulated kinase 1 and 2 (Erk1/2), where the binding of holotransferrin to TfR2 in a complex with Hfe induces Erk1/2 phosphorylation [17], and ii) the Hjv/BMP6R (bone morphogenetic protein-6 receptor) complex which activates Smad-signaling by forming the Smad1/5/8–Smad4 complex [18] (**Figure 2A**). Our study revealed a significant downregulation of miR-122 in the livers of Hjv^-/-^ animals (**Figure 2B**). Importantly, Hjv, like Hfe, is crucial for Hamp transcription maintenance [19] and the liver of Hjv^-/-^ mice exhibits iron overload [20]. Additionally, we measured miR-122 expression in the livers of transmembrane serine protease 6 (Tmprss6)^-/-^ mice. Tmprss6, a member of the type II transmembrane serine protease family, controls hepcidin transcription and inhibits this pathway by mediating Hjv proteolysis [21]. Notably, mutations preventing Tmprss6 activity lead to a rare inherited form of iron deficiency anemia called Iron-Refractory Iron Deficiency Anemia (IRIDA; [22,23]). Interestingly, our results identified a significant upregulation of miR-122 in the livers of Tmprss6^-/-^ transgenic mice (**Figure 2B**). These results support the conclusion that the iron sensing complex regulates miR-122 transcription downstream of Hjv/BMP6R signaling.

**Fig. 2.**
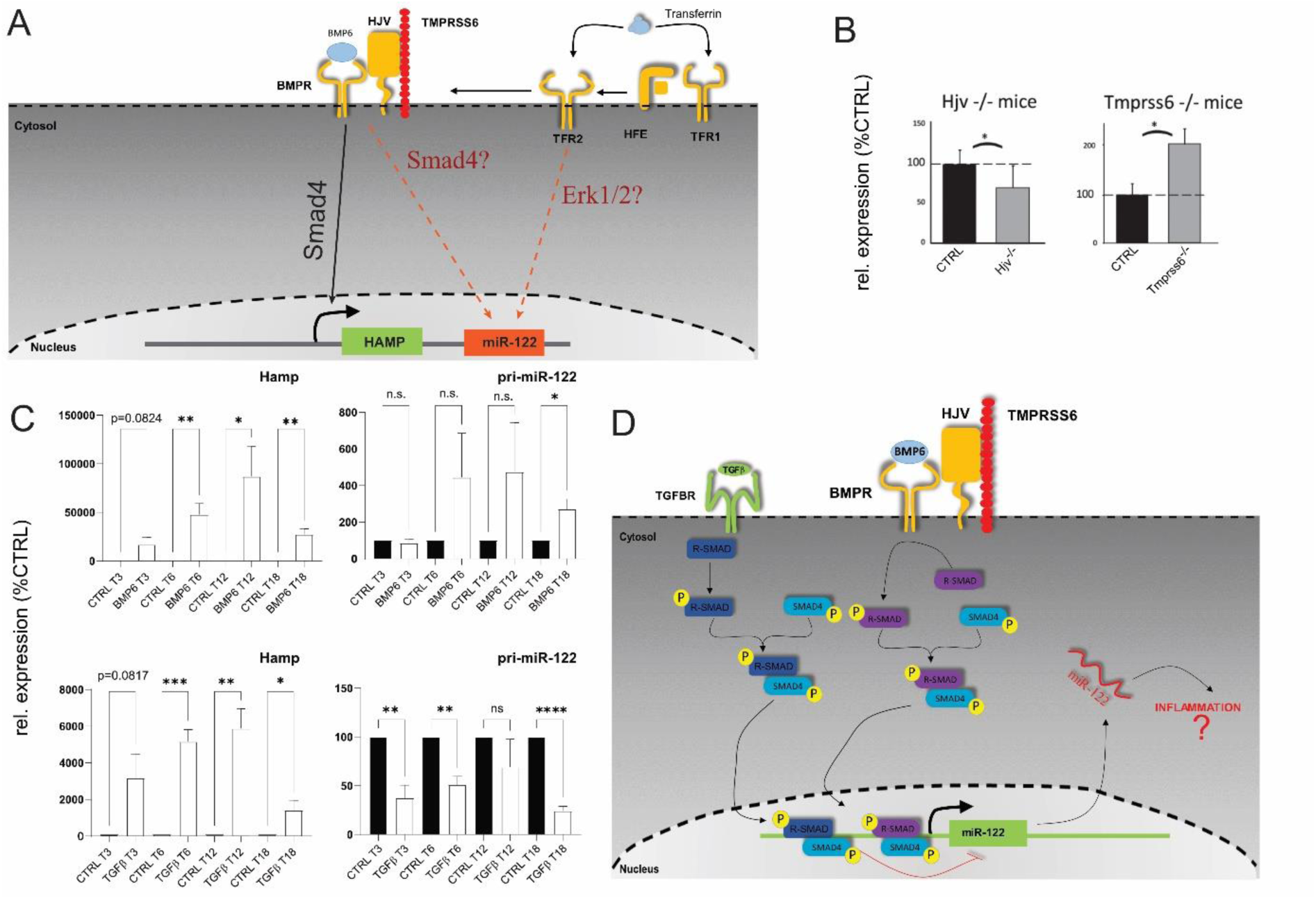
miR-122 is differentially regulated in in the liver of Hjv and Tmprss6 Ko animals. (**A**) Model of potential regulation of miR-122 associated with signaling downstream to Hfe. (**B**) qPCR analysis of miR-122 expression in the liver of Hjv KO (n = 5) and Tmprss6 KO animals (n = 5). (**C**) qPCR analysis of hepcidin and pri-miR-122 expression in primary mouse hepatocytes stimulated with either BMP6 (50 ng/mL; n = 4) or TGFβ1 (5 ng/mL; n = 4). (**D**) Proposed model of Smad-signaling upstream to miR-122 transcription. Results are represented as mean ± SD. Differences were evaluated by using t-test (ns = Not significant, *, p ≤ 0.05; **, p ≤ 0.01; ***, p ≤ 0.001).

### *MIR122* transcription is differentially regulated by TGFβ1 and BMP6

The observed alterations in miR-122 expression in the livers of Tmprss6^-/-^ and Hjv^-/-^ mice (**Figure 2A**), are consistent with the involvement of the Smad signaling pathway as an upstream regulator of miR-122 transcription. Notably, the expression of hepcidin, a key player in the response to inflammation, was shown to be also activated by transforming growth factor-beta 1 (TGFβ1; **Figure 2C**; [24,25]). Therefore, we investigated how miR-122 responded to TGFβ1 and BMP6 administration by treating murine primary hepatocytes with either TGFβ1 or BMP6. Although no significant changes in the expression of mature miR-122 were observed, analysis of the miR-122 primary transcript revealed distinct effects of TGFβ1 and BMP6 on *MIR122* transcription. TGFβ1 was found to inhibit *MIR122* transcription, while BMP6 significantly increased pri-miR-122 levels (**Figure 2D**). The efficacy of the treatment was confirmed by the expected upregulation of hepcidin mRNA in response to both TGFβ1 and BMP6 administration. Although the exact mechanism driving the transcriptional inhibition of *MIR122* in response to TGFβ1 administration is yet unknown, it can be speculated that this could be partially mediated via the activity of the Inhibitory Smads (I-Smads; [26]) as supported by the findings that TGFβ1, but not BMP6, upregulated the mRNA levels of several I-Smads, including Smad7 and the Smad-transcriptional co-repressors SnoN-1 and SnoN-2 (**Supplementary Figure S2A**; [27,28]).

### Activity of mouse *MIR122* promoter is increased by BMP6 and reduced by TGFβ1

The core promoter of miR-122 has been previously characterized in both humans and mice, unveiling its functional regulation by liver-enriched transcription factors (LEFT; [29,30]). In order to gain insight into the regulation of miR-122 transcription by cytokines, we employed the Evolutionary Conserved Regions (ECR) browser [31] to identify conserved regions upstream of the mouse miR-122 core promoter among different mammal species including *Mus musculus*, *Rattus norvegicus*, and *Homo sapiens sapiens* (**Supplementary Figure S3**). Four constructs were generated by cloning different length of the conserved region upstream of *MIR122* in a promoter luciferase vector, and their basal activity was assessed in a Huh-7 human hepatoma cell line. Consistent with previous studies [29,30], we found that the mouse *MIR122* core promoter is essential for maintaining miR-122 transcription. Notably, constructs containing longer upstream sequences exhibited higher activity, suggesting the presence of yet uncharacterized responsive elements within these sequences (**Supplementary Figure S4**). Subsequently, we transfected the constructs into mouse primary hepatocytes treated with either TGFβ1 or BMP6 (**Figure 3A**). Significantly, we observed that administration of TGFβ1 markedly reduced the activity of the construct with (pGL4.1-1.6Kb, p = 0.001 and pGL4.1-0.26Kb, p = 0.0031) and without (pGL4.1-0.95Kb, p = 0.0323) the core promoter region. Conversely, BMP6 administration significantly increased the activity of the construct containing the shorter promoter region (pGL4.1-0.75Kb, p = 0.0587 and pGL4.1-0.26Kb, p = 0.0114). To determine whether canonical or non-canonical Smad signaling is involved in mediating the regulation of miR-122 transcription by TGFβ1 and BMP6, we transfected primary hepatocytes isolated from wild-type mice with anti-Smad4 siRNA or a scrambled control oligo (SCR). Subsequently, we monitored the effect of TGFβ1 and BMP6 administration on the gene of interest using qPCR. Confirmation of Smad signaling attenuation in siRNA-transfected cells can be inferred through the significant reduction in hepcidin and Smad7 activation in response to TGFβ1 stimulation. Importantly, TGFβ1 effect on *MIR122* was blocked in cells transfected with anti-Smad4-siRNA (**Figure 3B**). Conversely, no significant BMP6-mediated regulation of hepcidin or *MIR122* transcription was observed, possibly indicating that primary hepatocytes lose their ability to respond to BMP6 administration after several days in culture. These findings support the hypothesis that, in mouse, TGFβ1 affects miR-122 transcription via canonical Smad signaling.

**Fig. 3:**
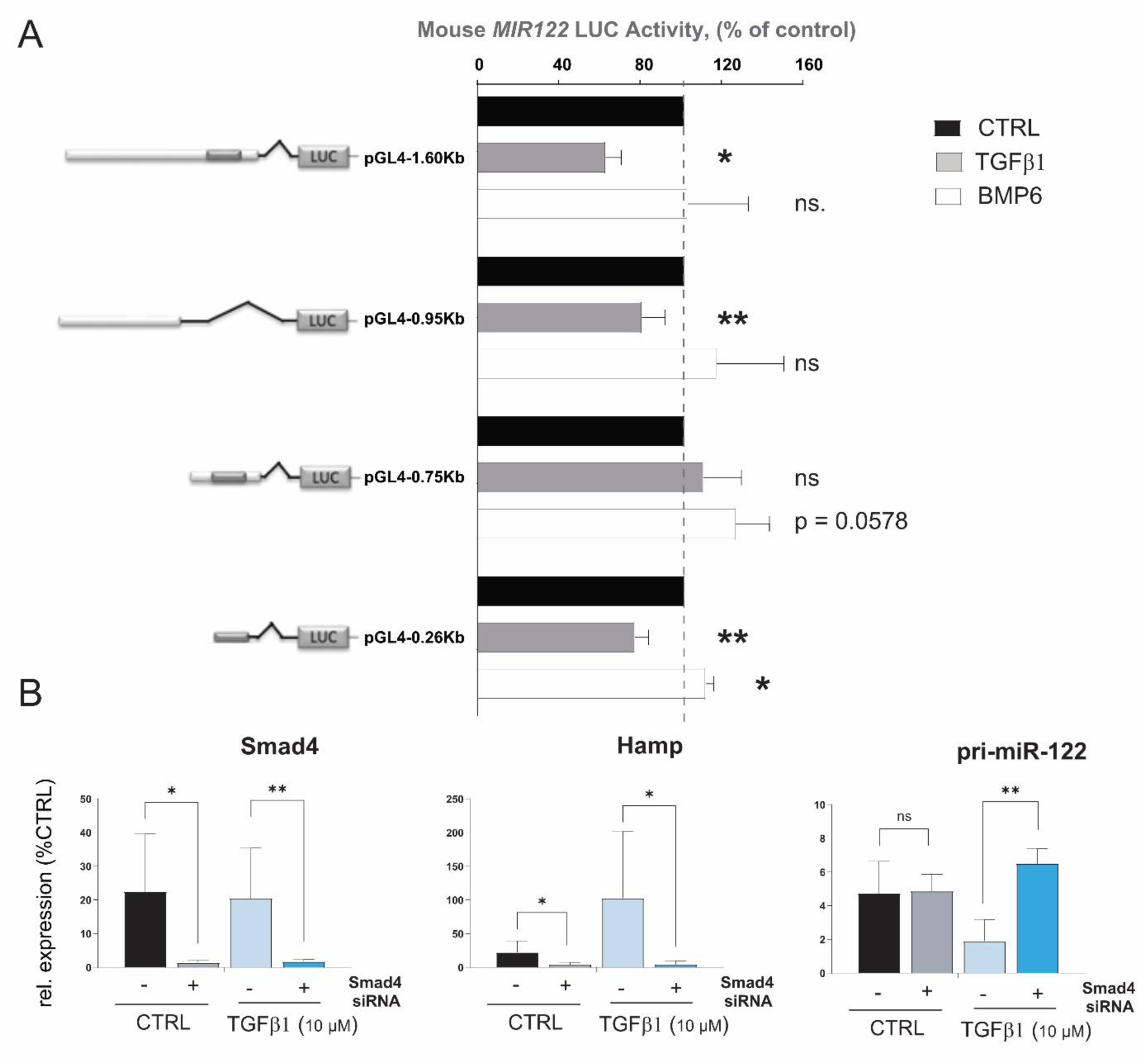
Mouse *MIR122* promoter regions respond to BMP6 and TGFβ stimulation. (**A**) Different lengths of upstream sequences of the mouse MIR122 promoter region were isolated and cloned in luciferase reporter vector. Hepa1-6 cells were transfected with constructs and treated with BMP6 or TGFβ1 or left untreated (n = 5). Cell were lysed 48 hours after the transfection and Luciferase activity measured. Data represent average luciferase activity in % of control (n = 5). (**B**) Mouse primary hepatocytes were transfected with anti Smad4 siRNA or scramble oligos (CTRL). Levels of pri-miR-122 were measured 18 hours after TGFβ1 administration (n = 4). Results are represented as mean ± SD, significant differences were evaluated by using One-way ANOVA or t-test as appropriate (ns = Not significant, *, p ≤ 0.05; **, p ≤ 0.01; ***, p ≤ 0.001).

### TGFβ1-mediated downregulation of miR-122 transcription is conserved in humans

To assess the impact of TGFβ1 and BMP6 on miR-122 expression in humans, we directly stimulated Huh-7 cells with these two cytokines and measured the expression of the genes of interest (GOIs) at different time points. Consistent with what was observed in mouse primary hepatocytes, no effect on mature miR-122 was observed (data not shown), while the expression of pri-miR-122 was significantly downregulated in response to TGFβ1 stimulation, showing a decrease at 3 (p = 0.004) and 6 hours (p = 0.003; **Figure 4A upper panel**). However, in contrast with the results in mouse, Huh-7 cells exhibited a significant reduction in pri-miR-122 levels upon BMP6 administration after 3 hours of incubation (p = 0.0111; **Figure 4A lower panel** and **Supplementary Figure S5**).

**Fig. 4.**
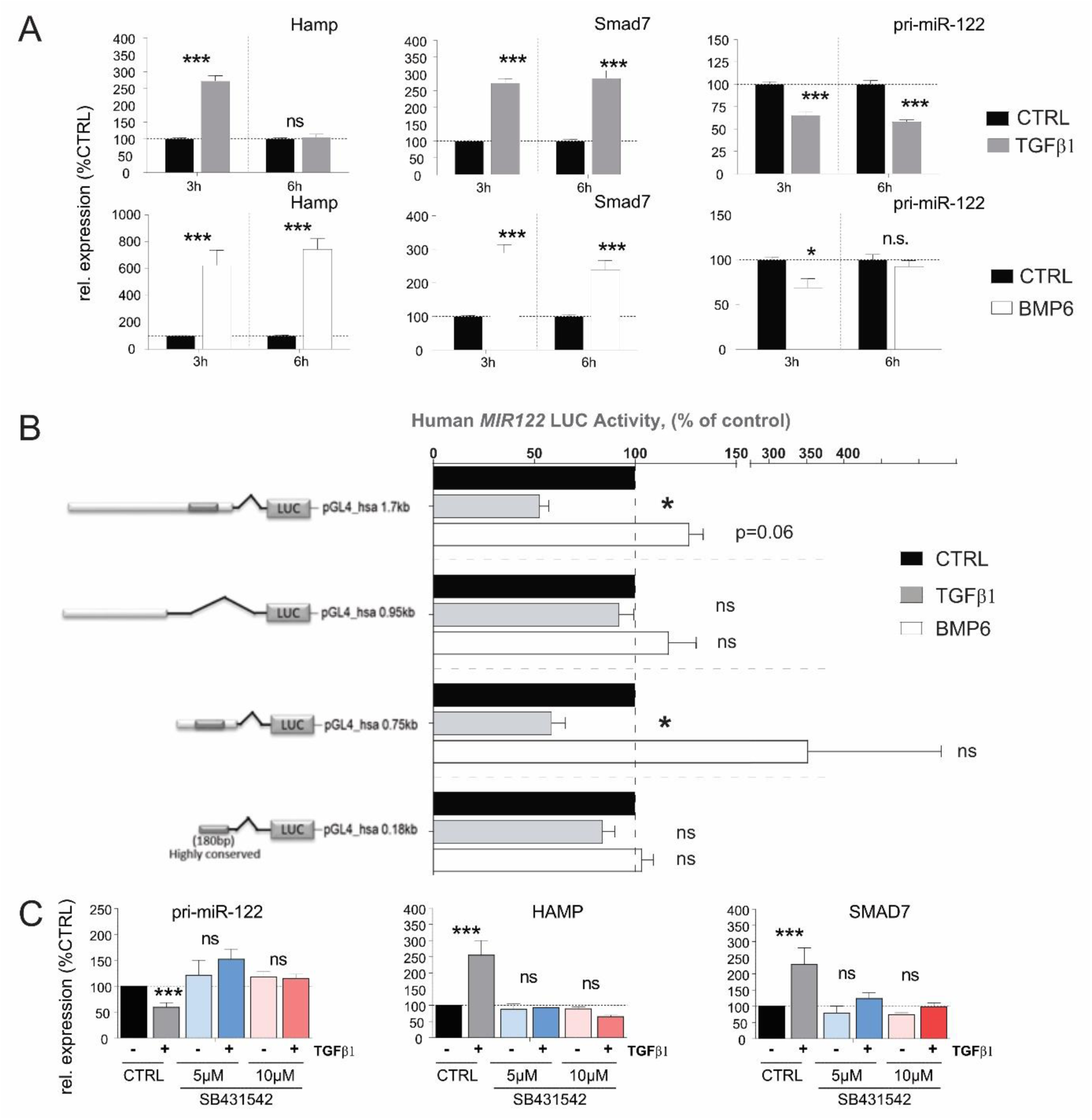
pri-miR-122 expression is differentially regulated in human hepatoma cells in response to TGFβ and BMP6 stimulation. (**A**) Effect of TGFβ1 and BMP6 stimulation on Hamp, Smad7 and pri-miR-122 expression in Huh-7 human-derived hepatoma cells, 3 and 6 hrs after stimulation (Normalized to GAPDH, n = 8). (**B**) Response of the human *MIR122* promoter to TGFβ1 and BMP6. Four promoter constructs encompassing different lengths of the human *MIR122* promoter were transfected into Huh-7 cells. After transfection, cells were starved for 24 hours and then treated with either TGFβ or BMP6. Data represent average luciferase activity in % of serum-free control (n = 5). (**C**) Serum starved Huh-7 cells were first incubated with TGFβ receptor type 1 inhibitor SB431542 (5 µM or 10 µM) for 3 h before stimulation with TGFβ1 or vehicle control for another 3 h. The levels of selected genes were measured by qPCR (Normalized to ACTB). Results are represented as mean ± SD, significant differences were evaluated by using One-way ANOVA or t-test as appropriate (ns = Not significant, *, p ≤ 0.05, ***, p ≤ 0.001).

To determine whether the observed discrepancy in BMP6 activity on human pri-miR-122 was species-specific or restricted to the Huh-7 cell line, we conducted an analysis of the human *MIR122* promoter region, which are highly conserved in mice and humans (**Supplementary Figure S3**). For this, four conserved regions upstream of the human *MIR122* promoter were cloned in the promoter-less pGL4.1-reporter vector (**Supplementary Figure S4, lower panel**) and the effect of TGFβ1 and BMP6 administration on the luciferase activity of these constructs was evaluated (**Figure 4B**). In accordance with what was observed in direct stimulation of Huh-7, our results showed that TGFβ1 significantly reduced the activity of the promoter constructs [pGL4_hsa-0.75Kb (p = 0.026) and the pGL4_hsa-1.7Kb (p = 0.009)]. In contrast to the findings in Huh-7 cells, BMP6 administration exhibited a trend towards increased activity of the pGL4_hsa-1.7Kb (p = 0.06) and of the pGL4_hsa-0.75Kb promoter constructs. This observed discrepancy suggests that either the activity of BMP6 is not conserved between humans and mice or that the *MIR122* promoter possesses additional BMP6 regulatory elements located further upstream or downstream of the *MIR122* core promoter that were not included in our constructs. Taken together, these data indicate that the inhibitory activity of TGFβ1 on miR-122 transcription is conserved in humans and mice, while the results obtained by BMP6 treatment on *MIR122* possibly indicates that the activity of this cytokine may not be conserved between the two species.

The TGFβ receptor complex transduces signals through what are known as canonical [32] and non-canonical [33] signaling pathways, hence to investigate which signaling pathway is required to mediate modulation of *MIR122* transcription, the TGF-Beta Type I receptor (TGFBR1) inhibitor SB431542 was administered to Huh-7 cells. It is important to note that TGFBR1 plays a crucial role in the canonical TGFβ signaling pathway [34]. Our results demonstrate that the administration of SB431542 effectively diminished both the inhibitory effect of TGFβ1 on *MIR122* transcription as well as blocked the transcriptional activation of HAMP and SMAD7 (**Figure 4C**). Consequently, we can conclude that TGFβ-mediated inhibition of miR-122 transcription is conserved and that this inhibitory effect is most likely mediated via canonical Smad signaling in both human and mouse.

### miR-122 is downregulated in the livers of LCMV infected mice and in CCl4 treated mice

To investigate the potential role of inflammation in the downregulation of miR-122 in liver diseases, we conducted a study using mice infected with Lymphocytic choriomeningitis virus (LCMV). LCMV belongs to the Arenaviridae family and is a non-lytic, single-stranded RNA virus. In our study, we focused on the LCMV WE strain, which induces acute hepatitis in mice that is subsequently cleared from the liver by CD8+ T cells within approximately two weeks [35]. To assess whether miR-122 expression is affected by viral infection in vivo, we infected wild-type mice (C57BL/6J) with LCMV WE viruses (2×10^6^ PFUs), and mice were sacrificed at various time points: before infection (day 0) and at 4, 6, 8, 10, 12, and 15 days post-infection (**Figure 5A**). As indicators of hepatocellular necrosis, we measured the levels of alanine aminotransferase (ALT) and bilirubin (**Figure 5B**). Interestingly, we observed a reduction in miR-122 expression in the liver of infected animals starting from day 6 post-infection (**Figure 5B**). Interestingly, increased levels of circulating cell-free miR-122 were simultaneously observed in the sera of these mice. Furthermore, we explored the effects of carbon tetrachloride (CCl4) administration, which is known to induce inflammatory and fibrotic signals, on miR-122 expression. We found that CCl4 treatment resulted in a significant downregulation of miR-122, even in the absence of hepatic cirrhosis or cancer (**Figure 5C**, **Figure 5D** and **Figure 5E**). Furthermore, we provided evidence that inflammatory and fibrotic signals associated with CCl4 administration lead to significant downregulation of miR-122 in the absence of liver cirrhosis or cancer. Collectively, these results support the conclusion that the downregulation of miR-122 observed in acute liver inflammation (e.g., LCMV WE infection) and chronic liver disease (e.g., CCl4 administration) is potentially a physiological response mediated by pro-inflammatory stimuli. These data also suggest the existence of a previously uncharacterized regulatory network linking infection and inflammation to declined miR-122 expression.

**Fig. 5.**
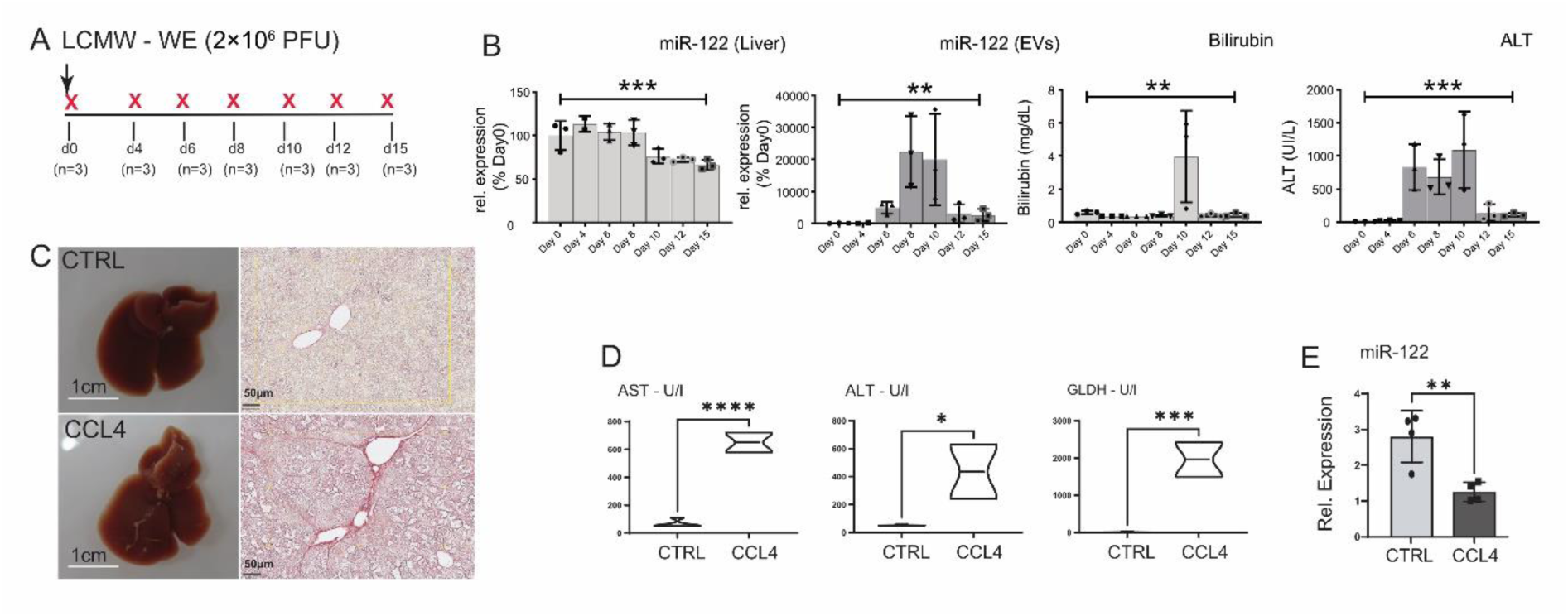
miR-122 expression is reduced in the liver of LCMV WE infected mice and in the livers of CCL4 treated animals. (**A**) Wild type C57-Bl6 mice were infected with 2×10^6^ PFUs of LCMV-WE and either euthanized before the infection (naïve) or at 4, 6, 8, 19, 12 and 15 days after infection (n = 3/time point). (**B**) qPCR analysis of miR-122 expression in RNA extracted from the liver and in the sera of animals included in the experiment shown in A; blood parameters for Bilirubin and ALT levels measure in the infected mice. (**C**) Representative macroscopical alterations and visualization of fibrosis by Sirius red staining mice that received CCL4 or vehicle (oil; CTRL) treatment for 8 weeks. (**D**) Blood parameters for AST, ALT and GLDH and (**E**) qPCR analysis of miR-122 levels measure in CCL4 and control mice. Results are represented as mean ± SD, significant differences were evaluated by using One-way ANOVA or t-test as appropriate (*, p ≤ 0.05; **, p ≤ 0.01; ***, p ≤ 0.001).

### Integrative analysis of gene expression in inflammation-driven liver tumors

To identify potential mediators of miR-122 activity in inflammation-driven liver cancer, CEL files from Affimetrix microarray included in our previous study [14] for the AlbLTα/β and Tet-O-Myc transgenic mice (accession number GSE102418) were reanalyzed with AltAnalyze [36] with respect to the identification of significantly upregulated genes in the liver of these animal models. Our analysis identified 60 genes that were significantly upregulated in the tumor compared to both near-tumor liver (NTL) and healthy tissue (**Figure 6A**). Out of these genes, 23 were predicted as potential targets of miR-122 according to the miRNA-target prediction database *miRWalk* (**Figure 6B**; [37]). To gain insight into the potential function of these genes, we performed an enrichment analysis using ShinyGO [38], which suggested that these genes are potentially involved in controlling the G2/M transition of the mitotic cell cycle (**Figure 6C**). To independently validate our findings, five genes were selected based on their expression and folds of upregulation in the tumor tissue [i.e., Anillin (Anln); Hyaluronan Mediated Motility Receptor (Hmmr); Targeting protein for Xklp2 (Tpx2); Cyclin B1 (Ccnb1) and Cyclin B2 (Ccnb2)] and their expressions were measured in liver tissue of various mouse models, including the livers of TRAF2/Casp-8^LPC-KO^ (**Figure 6D**), in the livers of mice overexpressing the oncogenes c-Myc and Neuroblastoma RAS viral oncogene homolog (NRAS; **Figure 6E** and **6F**), as well as in the livers of various transgenic mouse lines that spontaneously developed liver cancer (i.e., Nemo/Ripk3, Tak1^lpc-ko^, Ikk2/Casp8 and Traf2/Ripk1 ^lpc-ko^; **Supplementary Figure S6**; [14]). Importantly, the expression of the selected GOIs was consistently upregulated in diseased liver tissue of the animal models analyzed, suggesting their potential involvement in liver cancer development.

**Fig. 6.**
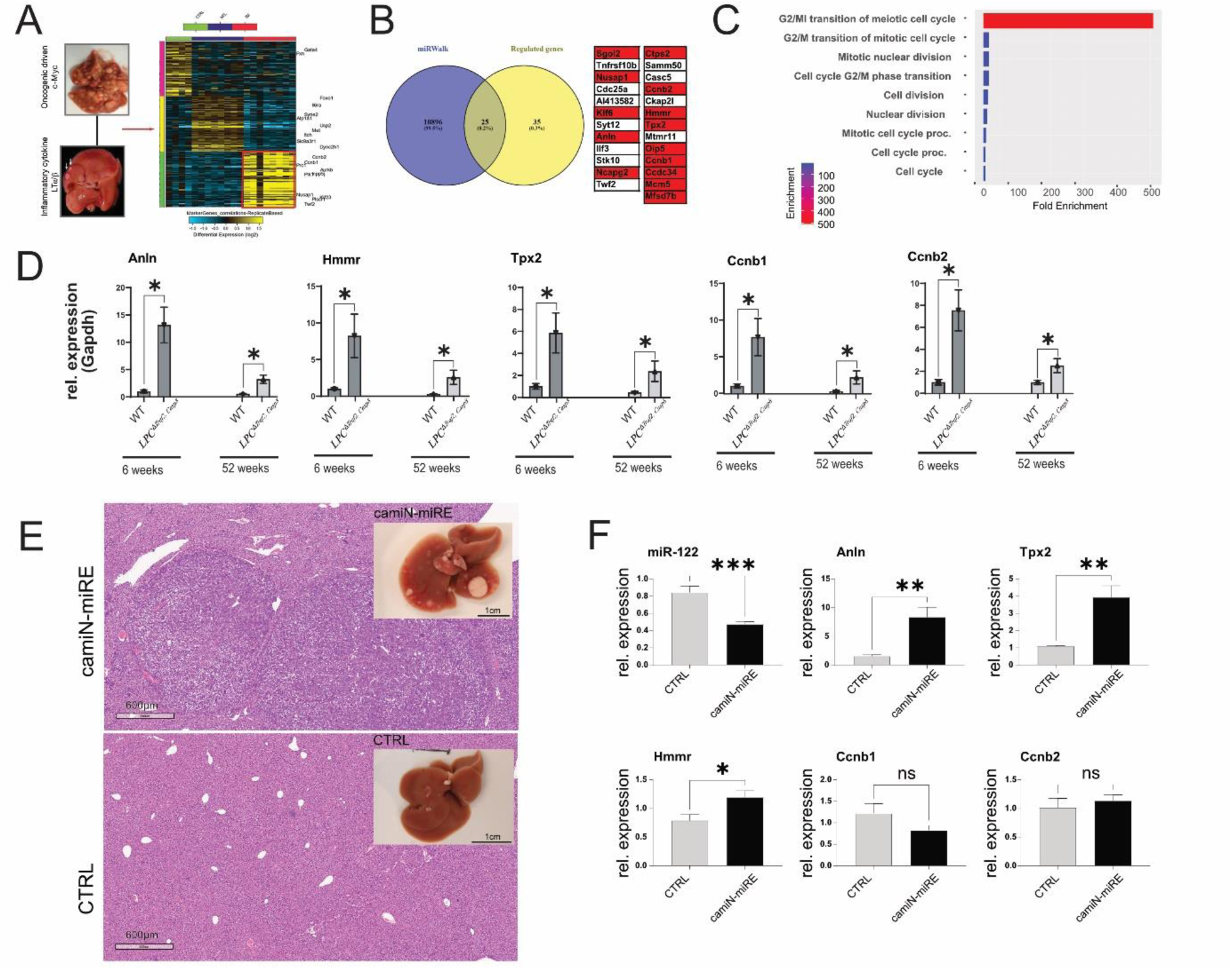
Analysis of gene expression in the livers of LT and c-myc mice. (**A**) Unsupervised hierarchical clustering of the genes regulated in the livers of LT α/β and c-Myc mice overexpressing mice. The color scale illustrates the fold change of mRNAs across the samples (Generated by Altanalyze). (**B**) Overlap between significantly downregulated genes and miRWalk predicted miR-122 targets. (**C**) ShinyGO enrichment analysis of potential miR-122 targets (**D**) qPCR analysis of the expression of highly enriched genes in liver of LPC^Δ*Traf2*^, *^Casp8^* animals at 6 and 52 weeks (n = 4). (**E**) Representative livers and HE staining of camiN-mirE and matching control (CTRL) untreated mice. (**F**) qPCR analysis of miR-122, Anln, Hmmr, Tpx2, Ccnb1, and Ccnb2, in the RNAs extracted from the livers of camiN-mirE (n =7) and CTRL (n = 4) mice. Results are represented as mean ± SD, significant differences were evaluated by using t-test (ns = Not significant; *, p ≤ 0.05; **, p ≤ 0.01; ***, p ≤ 0.001).

### Analysis of Anln, Hmmr, Tpx2, Ccnb1, and Ccnb2 expression in CCl4 injected mice and in activated HSCs

The data presented indicate that Anln, Hmmr, Tpx2, Ccnb1, and Ccnb2 may be influenced by changes in miR-122 expression and potentially contribute to liver cancer development. Our analysis also identified a significant downregulation of miR-122 in the livers of CCl4-injected mice (**Figure 5**). In order to investigate the potential involvement of Anln, Hmmr, Tpx2, Ccnb1 and Ccnb2 in the fibrotic process, the expressions of these genes were analyzed in the livers of CCl4-treated mice and found to be significantly upregulated (**Figure 7A**). Subsequently, the expression of Anln, Hmmr, Tpx2, Ccnb1 and Ccnb2 was measured in activated hepatic stellate cells (HSCs), which have been shown to become myofibroblast-like cells known to contribute to the progression of liver fibrosis through the deposition of extracellular matrix proteins [39]. Specifically GOIs’ expression was measured in the RNAs isolated from freshly isolated (day 0) and in activated (day 7) HSCs (**Figure 7B**). Our analysis revealed a significant increase in the expression of Anln, Hmmr, and Tpx2 in culture-activated HSCs, while the expression of Ccnb1 was downregulated and Ccnb2 remained unchanged. Taken together, these results suggest that upregulation of Anln, Hmmr, and Tpx2, but not of Ccnb1 and CCnb2, could potentially be involved in the process of fibrogenesis and activation of HSCs.

### Analysis of miR-122 responsive genes in GEO, TCGA and in Huh-7 human hepatoma cell

The downregulation of miR-122 in chronic liver diseases was further supported by the analysis of available databases, including GEO (e.g., accession number GSE51429, **Supplementary Figure S7**) and The Cancer Genome Atlas (TCGA; **Supplementary Figure S8**). To this end, detailed analysis of the expression data from the TCGA cohort of patients with hepatocellular carcinoma (LIHC) shows that miR-122 was significantly downregulated already at the early stages of HCC, while the Kaplan-Meier curve shows that patients with higher miR-122 expression have a significantly higher probability of survival than patients with lower miR-122 expression (p = 0.014; **Supplementary Figure S8**). Notably, and in agreement with what was observed in the mouse models of liver cancer, ANLN, HMMR, TPX2, CCNB1, and CCNB2 expressions were found to be significantly upregulated in TCGA-LIHC (**Supplementary Figure S9A**). Furthermore, the Kaplan-Meier curves for ANLN, HMMR, TPX2, CCNB1 and CCNB2 showed that a high expression of these genes was associated with an unfavorable prognosis (**Figure 7A** and **Supplementary Figure S9B**), which corroborated the analysis of ANLN, HMMR, TPX2, CCNB1 and CCNB2 in publicly available dataset from GEO (e.g., GSE62232, **Supplementary Figure S7** and GSE6764, **Supplementary Figure S10**). To investigate the potential (direct or indirect) regulation of ANLN, HMMR, TPX2, CCNB1, and CCNB2 expression by miR-122, human hepatoma cells Huh-7 were transfected with an antagomiR targeting miR-122 [10], and the expression of these genes was evaluated by qPCR. Our findings demonstrate that RNA levels of ANLN, HMMR, TPX2, CCNB1, and CCNB2 significantly increased in response to antagomir-mediated inhibition of miR-122 (**Figure 7B**). This supports the conclusion that miR-122 plays a role in controlling the expression of the aforementioned genes in Huh-7 cells, either directly or indirectly.

**Fig. 7.**
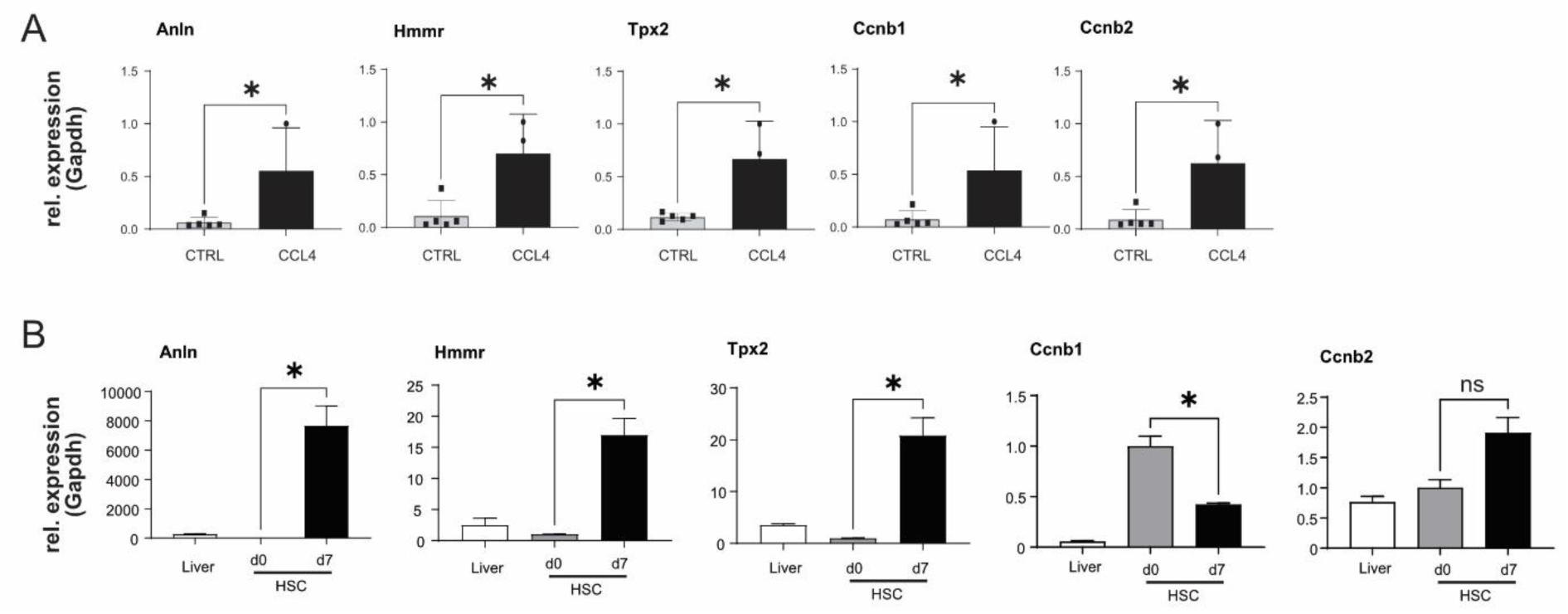
Analysis of gene expression in the livers of CCL4 injected mice and in activated HSCs. (**A**) qPCR expression of Anln, Hmmr, Tpx2, Ccnb1, and Ccnb2, in the livers of CCL4-treated (n =5) and oil injected control (CTRL) mice (n = 5). (**B**) Relative expression of GOIs in naïve and activated mouse HSCs and in the RNA extracted from the livers WT animals (n = 3). Results are represented as mean ± SD, significant differences were evaluated by using t-test (ns = Not significant; *, p ≤ 0.05).

## Discussion

Persistent inflammation has emerged as a significant contributor to the development and progression of malignancies, including hepatocellular carcinoma [1,2]. HCC is strongly associated with hepatic injury and inflammation, as more than 90% of HCC cases arise in the context of such conditions. Consequently, comprehending the underlying inflammatory signaling pathways in chronic liver diseases becomes crucial for identifying prognostic and diagnostic biomarkers as well as potential targets for stratification and treatment. This study employed an integrative approach to identify global changes in miRNA and mRNA expression that could potentially be involved in non-resolving inflammation-associated hepatocarcinogenesis. By applying high-throughput analysis of miRNAs in animal models of inflammation-driven liver cancer such as AlbLTα/β and Tet-O-Myc transgenic mice, we identified miR-122 as the most downregulated miRNA in tumor tissue. Although miR-122 is well-characterized, the mechanisms responsible for its decline in chronic liver diseases are not yet fully understood.

The present study made an important discovery regarding the transcriptional regulation of miR-122. We found that Tmprss6 and Hjv, which are upstream transcriptional regulators of the peptide hormone hepcidin, also play a role in the transcriptional regulation of miR-122. Overall, these data support the conclusion that persistent inflammation and HCC development involve complex interplay between miRNA expression, iron homeostasis, and innate immunity. Understanding these relationships may provide valuable insights for the identification of novel biomarkers, therapeutic targets, and potentially in the improvement of patient stratification and treatment strategies. Furthermore, our study demonstrates that BMP6 and TGFβ1, cytokines known to transactivate hepcidin transcription, exert differential regulation on the mouse promoter of *MIR122*. Importantly, we observed the conservation of TGFβ1 activity not only between human and mouse but also in rat (**Supplementary Figure S11**). While previous research has indicated that miR-122 influences cytokine-mediated signaling pathways [40], to the best of our knowledge our study is the first to establish a connection between inflammation, iron homeostasis, and the TGFβ-family members BMP6 and TGFβ1 to the control of miR-122 transcription.

Based on the findings presented in this study, we propose that the expression of miR-122 in the liver is governed by at least two locally competing regulatory loops. Firstly, a positive feedback loop (**Figure 8, model 1**) is established, wherein liver sinusoidal endothelial cells (LSEC) secrete BMP6 [41] to co-regulate Hamp and miR-122 expressions in response to changes in body iron requirements. Secondly, a negative feedback loop (**Figure 8, model 2**) is activated in response to local injury or inflammation, whereby TGFβ1 secreted by activated HSCs or immune cells suppresses *MIR122* transcription to trigger inflammatory responses. This hypothesis gains support from the notable observation that TGFβ1 administration significantly reduces Hjv expression in mouse hepatocytes (**Supplementary Figure S2B**). This may serve as an alternative mechanism to ensure the deactivation of iron-mediated modulation of miR-122. Notably, Yin et al. have previously demonstrated that miR-122 targets the TGFβ1 signaling pathway in both humans and mice [42]; This, along with the outcomes obtained in our study, leads us to suggest that TGFβ1 and miR-122 mutually regulate each other through a negative feedback loop. Notably, dysregulation of this regulatory loop may play a pivotal role in the development and progression of hepatocellular carcinoma. Literature mining has further revealed an inverse correlation between miR-122 and TGFβ1 levels in patients with liver diseases, including non-alcoholic fatty liver disease [43], primary biliary cholangitis [44], HCC [45], and viral hepatitis [46].

**Fig. 8.**
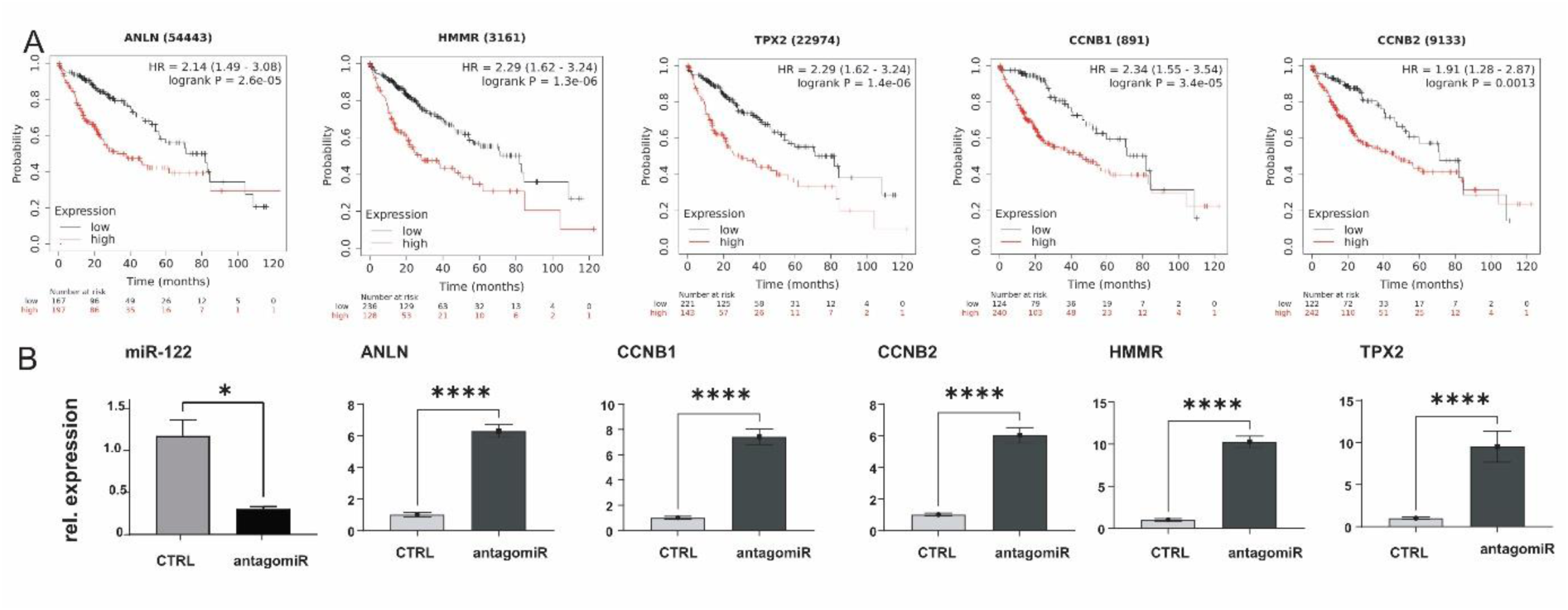
ANLN, HMMR, TPX2 CCNB1 and CCNB2, are significantly upregulated in human liver cancer. (**A**) Kaplan-Meier survival curves for ANLN, HMMR, TPX2, CCNB1 and CCNB2, in TCGA LIHC cohorts. (**B**) qPCR expression for ANLN, HMMR, TPX2, CCNB1 and CCNB2 in RNAs extracted from Huh-7 hepatoma cells transfected with either control scrambled oligo or miR-122 antagomiR (n = 5). Results are represented as mean ± SD, significant differences were evaluated by using t-test (*, p ≤ 0.05; ****, p ≤ 0.0001).

**Fig. 9.**
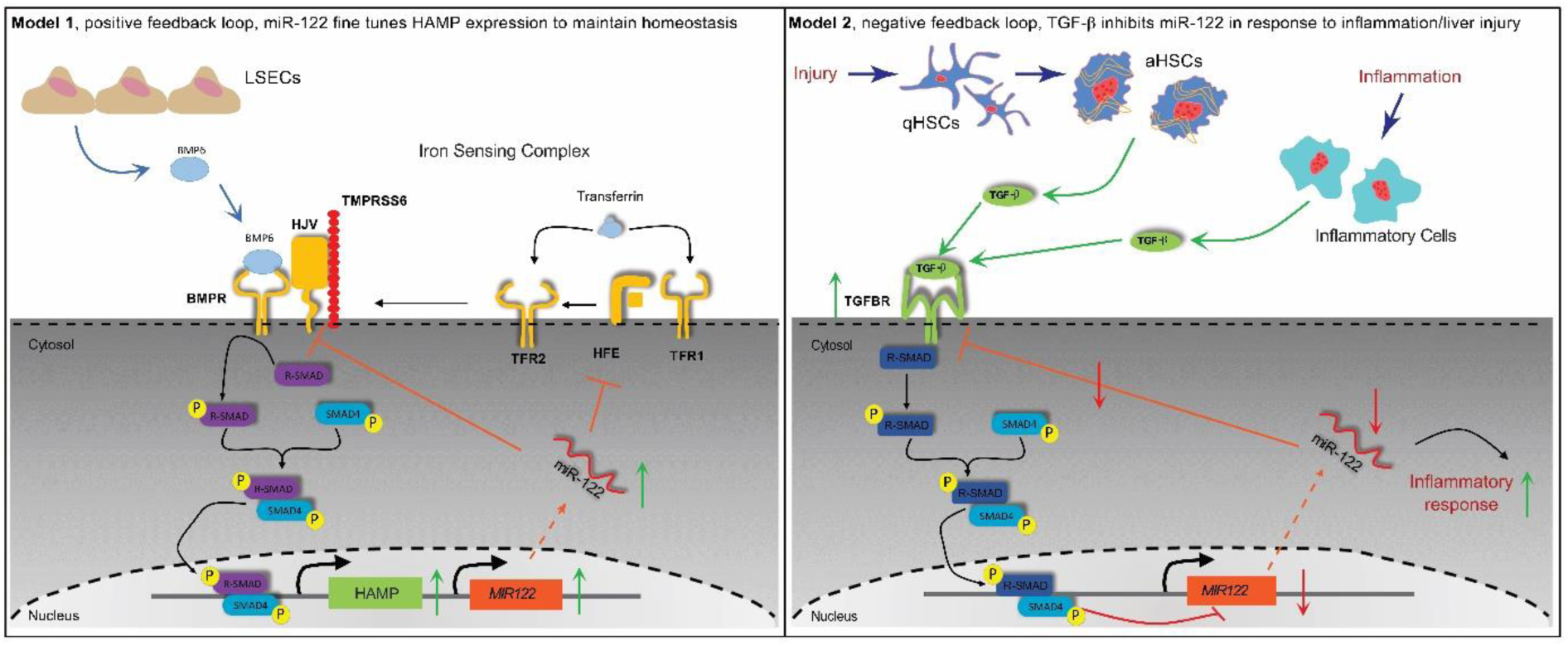
Proposed models for regulatory networks up-steam to miR-122. (**A**) **Model 1**; positive feedback loop; during normal control of iron homeostasis. LSEC release BMP6 in response to high iron storage, BMP6 activated Smad-signaling leading to both the activation of HAMP and miR-122 transcription. In this model the role of miR-122 is to fine-tune HAMP transcription via inhibiting HJV and HFE translation. (**B**) **Model 2**; negative feedback loop, in response to liver injury (e.g., HSCs activation) and inflammation. Activate HSCs or Immune cells release TGFβ1. Following TGFβ1 binding to its receptor Canonical Smad-signaling is activated leading to i) downregulation of *MIR122* transcription; and ii) Increase of TGF-signaling thought the reduction of miR-122-mediated inhibition. In this model, the role of miR-122 down-regulation is activate inflammatory response via release of miR-122 inhibitions on its target genes.

Hence, liver injury lead to an upregulation of TGFβ1, which plays a critical role in liver regeneration [47,48] and terminates the proliferation of hepatocytes [49], that in turn, can cause a decrease in the transcription of *MIR122* in chronic injured liver. Our study emphasizes that abnormally low levels of miR-122 could trigger a negative feedback loop, contributing to persistent inflammation, increased susceptibility to chronic liver disease, and elevated risk of HCC development. It is important to note that while we did not observe significant changes in mature miR-122 levels, previous studies by Gatfield and colleagues have reported similar inconsistencies between miR-122 and pri-miR-122. They demonstrated that pri-miR-122 transcription follows a circadian rhythm, whereas mature miR-122 remains relatively stable throughout the day [50,51]. One possible explanation is that once miR-122 binds to its target mRNA, they remain stably associated with RISC complexes. Therefore, it has been speculated that the synthesis of new miR-122 molecules, rather than the overall cellular levels, may be the critical determinant of miRNA-mediated gene silencing efficiency.

Our study has identified several potential miR-122-responsive genes in livers of animals with inflammation-associated liver cancer, among them ANLN, TPX2, HMMR, CCNB1, and CCNB2. Through enrichment analysis and literature mining, we have gained insights suggesting the potential involvement of these genes in microtubule and cytoskeletal dynamics, as well as the mitotic checkpoint, indicating a novel potential role for miR-122 in controlling cellular growth and migration. Furthermore, our research shows that the regulatory control potentially exerted by miR-122 on these genes in mice is probably conserved in humans, although at the current stage of our research we cannot determine whether miR-122 exerts its control over these genes through direct binding to miRNA responsive elements (MREs) present in GOIs 3UTRs or indirectly through the modulation of yet unknown regulators such as specific signaling pathways or transcription factors. On the basis of our results, we propose that excessive activation of regulatory networks associated with cytokine-mediated deregulation of miR-122 expression may contribute to the transition of acute, nonresolving inflammation into full-scale chronic liver disease.

## Material and Methods

### Human and mouse *MIR122* promoter cloning

Mouse and human *MIR122* promoter regions were amplified from genomic DNA and cloned in the pGL4.10[luc2] promoter-less luciferase plasmid (Promega, Walldorf, Germany,). Restriction sites for NheI, HindIII for cloning into the selected sites of the reporter vector were included to the 5’ end of the respective primers (**Supplementary Table S1**). Following PCR amplification, amplicons were subjected to agarose gel electrophoresis for product size selection and purified with QIAquick Gel Extraction Kit (Qiagen, Hilden, Germany). Ligation was performed as previously described [10]. The isolated plasmids were sequenced by the Genomics and Transcriptomics Laboratory (GTL, Heinrich Heine University, Düsseldorf) by means of Sanger-Sequencing.

### RNA isolation, quantification, cDNA synthesis, and qPCR analysis

RNA isolation, quantification, and reverse transcription were performed as previously described [10]. miRNA profiling was performed by using the miQPCR method [52]. miQPCR performs the universal reverse transcription of all the miRNAs contained in the RNA sample enabling the analysis of up to 100 individual miRNAs from every synthesized cDNA. qPCR reactions were carried out on a ViaA7 cycler (Thermo Fisher Scientific, Meerbusch, Germany) and amplicons were detected using SYBR Green I (GO-Taq PCR Master mix, Cat: A6002; Promega, Walldorf, Germany). Sequences of the primers used in this study are listed in **Supplementary Table S2**. Data analysis was carried out by using qBase [53]. Suitable reference genes were identified by using geNorm [54], and samples were normalized by using ΔΔCt [55]. For the experiments where no suitable reference genes were found, qPCR data were median normalized from within qBase by using a microarray-like approach as described [56]. GraphPad Prism was used to perform statistical analyses. Unpaired T-test of control group versus individual treatments or, when comparing more than two groups, One-Way analysis of variance (ANOVA) were carried out.

### Preparation of primary hepatocytes, HSCs and transfections of reporter vectors

HSCs were isolated from livers as previously described [47]. Primary murine hepatocytes were isolated according to an optimized protocol described previously [57], whereas primary rat hepatocytes were prepared as published elsewhere [52]. Plasmid carrying the human *MIR122* promoter regions were co-transfected in Huh-7 cells with the pRL-SV40 renilla vector for normalization, while plasmid carrying the mouse *MIR122* promoter regions were co-transfected in mouse primary hepatocytes. Huh-7 were transfected with linear 25 kDa polyethyleneimine (PEI, Polysciences Inc., Warrington, PA, USA) by using a 3:1 ratio of PEI/DNA (as described in [58]). Mouse primary hepatocytes were transfected by using RNAiMAX (Invitrogen, Waltham, MA, USA) by following the included instructions.

Following transfection, Promegás Dual-Luciferase Reporter Assay (Promega, Walldorf, Germany) was conducted in accordance with the manufactureŕs instructions. The chemiluminescence assay was performed in white opaque 96-well plates with 50 µL cell lysate and 50 µL of both, LARII reagent (Firefly luciferase substrate) and Stop and Glo reagent (Renilla luciferase substrate), in a GloMax Multi Plus Multiplate Reader (Promega, Walldorf, Germany) according to the preset protocol with an integrity time of 10 seconds for each read. Data were normalized by calculating ratios of Firefly/ Renilla activities to correct for possible variations in transfection efficiencies.

### Transfection and cytokine administration

For cytokine treatment and transfection, cells were seeded in 6 well plates 1 day before treatment and incubated in serum free medium. Transfection of siRNA and miRNA inhibitors were carried out by using Lipofectamine RNAiMax (Thermo Fisher Scientific, Meerbusch, Germany) in accordance with the manufactureŕs instructions. miR-122 inhibition was carried out by transfection with AntagomiR 122 (Miravirsen, SPC-3649; Roche, formerly Santaris Pharma). For Smad4 silencing in mouse primary hepatocytes, the mouse specific Smad4 siRNA (Cat.12791; Cell Signaling) and the Signal Silence Control siRNA (Cat.6568S; Cell Signaling, Danvers, MA, USA) were used. The day of the transfection cells were added of fresh medium (DMEM medium, 10% FCS and 1% penicillin/streptomycin) and transfected with either 100 nM of miR-122 antagomiR/ Smad4-siRNA or 100 nM of the corresponding negative controls. After 48 hours transfected cells were lysed for downstream analyses. Primary hepatocytes or cell lines were seeded in 6 well plates and cultured or transfected according to the protocols indicated above. For cytokine treatment cells were stimulated with either 5 ng/mL of TGFβ1 or 50 ng/mL of BMP6 (both from PeproTech, Rocky Hill, NJ, US). Where indicated, cells were administrated with 10 µM of the TGFβ receptor-I inhibitor SB431542 (Axon Medchem LLC, Groningen, Netherlands) and incubated for 3 hours before addition of TGFβ1.

### Transgenic animals, hydrodynamic tail vein injection and CCl4 mice

The generation and characterization of the double [TNF receptor-associated factor 2 (Traf2) and Caspase-8 (Casp8)] and the triple [Traf2, Casp8 and Receptor-interacting serine/threonine-protein kinase 3(Ripk3); and the Traf2, Casp8 and Inhibitor of nuclear factor kappa-B kinase subunit beta (Ikbkb)] KO mice included in this study was published by our group in [16]. The hydrodynamic tail vein injections (HDTVi) were performed as described in [14]. In brief, 5 weeks old wild-type C57BL/6J male mice were ordered from Janvier Labs (Le Genest, Pays de la Loire, France), let recover for two weeks before HDTVi. For HDTVi 7 weeks old mice were anesthetized with and Isoflurane (Piramal, Andhra Pradesh, India) and vectors [i.e., 25 µg transposon vector (carrying c-Myc and NRAS oncogenes) and 5 µg SB13 transposase vector] were delivered in 10% v/w body weight of physiological solution (0.9% NaCl) in the mouse’s tail vein. Animals were euthanized 6 weeks after the procedure. For fibrosis induction, wild-type mice (C57BL/6J) were injected twice a week in the peritoneum with 50 µL of CCl4 (6 µL CCl4/10 g body weight/ Oil up to 50 µL), while control animals were injected with 50 µL of Oil.

### Analyses of data from the Genome Omnibus repository

The miRNA and mRNA profiling data in animal model of liver cancer have been retrieved from GEO [15] under the super-series accession number GSE102418. This super-series is composed of the following sub-series: GSE102416, which contains the raw data of the miRNAs expression analyses, and GSE102417 which contains the CEL files of the Affymetrix microarrays. Enrichment analysis was performed with ShinyGO [38] by uploading list of significantly upregulated genes, which were predicted to be miR-122 targets by *miRWalk* [37] target prediction database. Differentially regulated genes in the Affymetrix arrays were identified by using AltAnalyze [36] with fold changes cutoff ≥ 2; p ≤ 0.05.

### Analyses of The Cancer Genome Atlas (TCGA) data and Kaplan-Meier survival curves

Some of the result shown in this study are based upon data generated by the TCGA Research Network: https://www.cancer.gov/tcga. Kaplan-Meier survival curves of LIHC patients for miR-122, ANLN, TPX2, HMMR, CCNB1, and CCNB2 were generated by using the KM Plotter [59].

### Statistical Analysis and Imaging Software

Statistical analyses were carried out using GraphPad Prism (version 9.4.1). To evaluate whether samples were normally distributed, the D’Agostino and Pearson normality tests were carried out. When the sample distribution passed the normality test, then parametric tests were carried out (i.e., one-way analysis of variance/ANOVA for three or more samples and a two-tailed T-test for two samples). When the samples did not pass the normality test, non-parametric tests were applied (i.e., the Kruskal-Wallis test for three or more samples and the Mann-Whitney test for two samples). The data were considered significant at a p value ≤ 0.05. Images were prepared using Affinity Designer (version 1.10.4.1198).

## Abbreviations

miRNA, microRNA; MRE, miRNA responsive elements; HCC, hepatocellular carcinoma; NTL, near-tumor liver; Hamp, hepcidin; Hfe, hemochromatosis; Hjv, hemojuvelin; Tmprss6, transmembrane serine protease 6; HH, hereditary hemochromatosis; DEN, Diethylnitrosamine; Traf2, TNF receptor-associated factor 2; Casp-8, Caspase-8; Erk1/2, extracellular signal-regulated kinase 1 and 2; TGFβ, transforming growth factor-beta; TGFBR1, TGF-Beta Type I receptor; Bmp6r, bone morphogenetic protein-6 receptor; Bmp6, bone morphogenetic protein-6; IRIDA, Iron-Refractory Iron Deficiency Anemia; LCMV, Lymphocytic choriomeningitis virus; CCl4, carbon tetrachloride; ALT, alanine aminotransferase; HDTVi, Hydrodynamic tail vein injection; LIHC, Liver Hepatocellular Carcinoma; LEFT, liver enriched transcription factors; TCGA, The Cancer Genome Atlas; GEO, Genome Omnibus; GOIs, genes of interest; ECR browser, Evolutionary Conserved Regions browser; HSCs, hepatic stellate cells; LSEC, liver sinusoidal endothelial cells; NRAS, Neuroblastoma RAS viral oncogene homolog; Anln, Anillin; Hmmr, Hyaluronan Mediated Motility Receptor; Tpx2, Targeting protein for Xklp2; Ccnb1, Cyclin B1; Ccnb2, Cyclin B2.

## Author contributions

M.P., designed research, performed experiments and wrote the manuscript. C.K., provided rat parenchymal cells. M.V., provided the mouse models of liver cancer. V.B., prepared the CCl4 treated mice. H.X., P.S., and P.L., performed the experiments with the LCMV WE mice. C.R. and T.L., designed research. M.C., conceived, designed, performed experiments, and wrote the manuscript. All authors read and approved the manuscript.

## Ethics approval

The relevant official authority for animal protection (Landesamt für Natur-, Umwelt-und Verbraucherschutz, LANUV, Nordrhein-Westfalen, Recklinghausen, Germany) approved these studies. The HDTV injections (81-02.04.2020.A017), CCl4 administration (81-02.04.2020.A017) and LCMV WE infections were approved by the LANUV. Animals received care according to the German animal welfare act. Animal experiments included in this study were carried out in compliance with the Act No 246/1992 Coll. on the protection of animals against cruelty.

## Informed consent statement

Not applicable.

## Data availability statement

The datasets used and/or analyzed during the current study are available from the corresponding author on reasonable request.

## Acknowledgements

The authors are grateful to Claudia Rupprecht for the technical assistance. We are grateful to Martina Muckenthaler for providing the liver RNAs from Hjv^-/-^ mice and to Andrew J Ramsay for providing the RNA from the livers of Tmprss6^-/-^ mice.

## Reference

1. Zhao, H.; Wu, L.; Yan, G.; Chen, Y.; Zhou, M.; Wu, Y.; Li, Y. Inflammation and tumor progression: signaling pathways and targeted intervention. Signal Transduct Target Ther 2021, 6, 263, doi:10.1038/s41392-021-00658-5.

2. Yu, L.X.; Ling, Y.; Wang, H.Y. Role of nonresolving inflammation in hepatocellular carcinoma development and progression. NPJ Precis Oncol 2018, 2, 6, doi:10.1038/s41698-018-0048-z.

3. Vidigal, J.A.; Ventura, A. The biological functions of miRNAs: lessons from in vivo studies. Trends Cell Biol 2015, 25, 137–147, doi:10.1016/j.tcb.2014.11.004.

4. Halasz, T.; Horvath, G.; Par, G.; Werling, K.; Kiss, A.; Schaff, Z.; Lendvai, G. miR-122 negatively correlates with liver fibrosis as detected by histology and FibroScan. World J Gastroenterol 2015, 21, 7814–7823, doi:10.3748/wjg.v21.i25.7814.

5. Padgett, K.A.; Lan, R.Y.; Leung, P.C.; Lleo, A.; Dawson, K.; Pfeiff, J.; Mao, T.K.; Coppel, R.L.; Ansari, A.A.; Gershwin, M.E. Primary biliary cirrhosis is associated with altered hepatic microRNA expression. J Autoimmun 2009, 32, 246–253, doi:10.1016/j.jaut.2009.02.022.

6. Li, C.; Wang, Y.; Wang, S.; Wu, B.; Hao, J.; Fan, H.; Ju, Y.; Ding, Y.; Chen, L.; Chu, X.;, et al. Hepatitis B virus mRNA-mediated miR-122 inhibition upregulates PTTG1-binding protein, which promotes hepatocellular carcinoma tumor growth and cell invasion. J Virol 2013, 87, 2193–2205, doi:10.1128/JVI.02831-12.

7. Luna, J.M.; Scheel, T.K.; Danino, T.; Shaw, K.S.; Mele, A.; Fak, J.J.; Nishiuchi, E.; Takacs, C.N.; Catanese, M.T.; de Jong, Y.P.;, et al. Hepatitis C virus RNA functionally sequesters miR-122. Cell 2015, 160, 1099–1110, doi:10.1016/j.cell.2015.02.025.

8. Kutay, H.; Bai, S.; Datta, J.; Motiwala, T.; Pogribny, I.; Frankel, W.; Jacob, S.T.; Ghoshal, K. Downregulation of miR-122 in the rodent and human hepatocellular carcinomas. J Cell Biochem 2006, 99, 671–678, doi:10.1002/jcb.20982.

9. Hsu, S.H.; Wang, B.; Kota, J.; Yu, J.; Costinean, S.; Kutay, H.; Yu, L.; Bai, S.; La Perle, K.; Chivukula, R.R.;, et al. Essential metabolic, anti-inflammatory, and anti-tumorigenic functions of miR-122 in liver. J Clin Invest 2012, 122, 2871–2883, doi:10.1172/JCI63539.

10. Castoldi, M.; Vujic Spasic, M.; Altamura, S.; Elmen, J.; Lindow, M.; Kiss, J.; Stolte, J.; Sparla, R.; D’Alessandro, L.A.; Klingmuller, U.;, et al. The liver-specific microRNA miR-122 controls systemic iron homeostasis in mice. J Clin Invest 2011, 121, 1386–1396, doi:10.1172/JCI44883.

11. Muckenthaler, M.; Roy, C.N.; Custodio, A.O.; Minana, B.; deGraaf, J.; Montross, L.K.; Andrews, N.C.; Hentze, M.W. Regulatory defects in liver and intestine implicate abnormal hepcidin and Cybrd1 expression in mouse hemochromatosis. Nat Genet 2003, 34, 102–107, doi:10.1038/ng1152.

12. Papanikolaou, G.; Samuels, M.E.; Ludwig, E.H.; MacDonald, M.L.; Franchini, P.L.; Dube, M.P.; Andres, L.; MacFarlane, J.; Sakellaropoulos, N.; Politou, M.;, et al. Mutations in HFE2 cause iron overload in chromosome 1q-linked juvenile hemochromatosis. Nat Genet 2004, 36, 77–82, doi:10.1038/ng1274.

13. Michels, K.; Nemeth, E.; Ganz, T.; Mehrad, B. Hepcidin and Host Defense against Infectious Diseases. PLoS Pathog 2015, 11, e1004998, doi:10.1371/journal.ppat.1004998.

14. Roy, S.; Hooiveld, G.J.; Seehawer, M.; Caruso, S.; Heinzmann, F.; Schneider, A.T.; Frank, A.K.; Cardenas, D.V.; Sonntag, R.; Luedde, M.;, et al. microRNA 193a-5p Regulates Levels of Nucleolar-and Spindle-Associated Protein 1 to Suppress Hepatocarcinogenesis. Gastroenterology 2018, 155, 1951–1966 e1926, doi:10.1053/j.gastro.2018.08.032.

15. Patra, B.G.; Roberts, K.; Wu, H. A content-based dataset recommendation system for researchers-a case study on Gene Expression Omnibus (GEO) repository. Database (Oxford) 2020, 2020, 1, doi:10.1093/database/baaa064.

16. Vucur, M.; Ghallab, A.; Schneider, A.T.; Adili, A.; Cheng, M.; Castoldi, M.; Singer, M.T.; Buttner, V.; Keysberg, L.S.; Kusgens, L.;, et al. Sublethal necroptosis signaling promotes inflammation and liver cancer. Immunity 2023, doi:10.1016/j.immuni.2023.05.017.

17. Poli, M.; Luscieti, S.; Gandini, V.; Maccarinelli, F.; Finazzi, D.; Silvestri, L.; Roetto, A.; Arosio, P. Transferrin receptor 2 and HFE regulate furin expression via mitogen-activated protein kinase/extracellular signal-regulated kinase (MAPK/Erk) signaling. Implications for transferrin-dependent hepcidin regulation. Haematologica 2010, 95, 1832–1840, doi:10.3324/haematol.2010.027003.

18. Xia, Y.; Babitt, J.L.; Sidis, Y.; Chung, R.T.; Lin, H.Y. Hemojuvelin regulates hepcidin expression via a selective subset of BMP ligands and receptors independently of neogenin. Blood 2008, 111, 5195–5204, doi:10.1182/blood-2007-09-111567.

19. D’Alessio, F.; Hentze, M.W.; Muckenthaler, M.U. The hemochromatosis proteins HFE, TfR2, and HJV form a membrane-associated protein complex for hepcidin regulation. J Hepatol 2012, 57, 1052–1060, doi: 10.1016/j.jhep.2012.06.015.

20. Huang, F.W.; Pinkus, J.L.; Pinkus, G.S.; Fleming, M.D.; Andrews, N.C. A mouse model of juvenile hemochromatosis. J Clin Invest 2005, 115, 2187–2191, doi:10.1172/JCI25049.

21. Silvestri, L.; Pagani, A.; Nai, A.; De Domenico, I.; Kaplan, J.; Camaschella, C. The serine protease matriptase-2 (TMPRSS6) inhibits hepcidin activation by cleaving membrane hemojuvelin. Cell Metab 2008, 8, 502–511, doi:10.1016/j.cmet.2008.09.012.

22. Finberg, K.E.; Heeney, M.M.; Campagna, D.R.; Aydinok, Y.; Pearson, H.A.; Hartman, K.R.; Mayo, M.M.; Samuel, S.M.; Strouse, J.J.; Markianos, K.;, et al. Mutations in TMPRSS6 cause iron-refractory iron deficiency anemia (IRIDA). Nat Genet 2008, 40, 569–571, doi:10.1038/ng.130.

23. Krijt, J.; Fujikura, Y.; Ramsay, A.J.; Velasco, G.; Necas, E. Liver hemojuvelin protein levels in mice deficient in matriptase-2 (Tmprss6). Blood Cells Mol Dis 2011, 47, 133–137, doi:10.1016/j.bcmd.2011.04.009.

24. Chen, S.; Feng, T.; Vujic Spasic, M.; Altamura, S.; Breitkopf-Heinlein, K.; Altenoder, J.; Weiss, T.S.; Dooley, S.; Muckenthaler, M.U. Transforming Growth Factor beta1 (TGF-beta1) Activates Hepcidin mRNA Expression in Hepatocytes. J Biol Chem 2016, 291, 13160–13174, doi:10.1074/jbc.M115.691543.

25. Canali, S.; Vecchi, C.; Garuti, C.; Montosi, G.; Babitt, J.L.; Pietrangelo, A. The SMAD Pathway Is Required for Hepcidin Response During Endoplasmic Reticulum Stress. Endocrinology 2016, 157, 3935–3945, doi:10.1210/en.2016-1258.

26. Hata, A.; Chen, Y.G. TGF-beta Signaling from Receptors to Smads. Cold Spring Harb Perspect Biol 2016, 8, doi:10.1101/cshperspect.a022061.

27. Kautz, L.; Meynard, D.; Monnier, A.; Darnaud, V.; Bouvet, R.; Wang, R.H.; Deng, C.; Vaulont, S.; Mosser, J.; Coppin, H.;, et al. Iron regulates phosphorylation of Smad1/5/8 and gene expression of Bmp6, Smad7, Id1, and Atoh8 in the mouse liver. Blood 2008, 112, 1503–1509, doi:10.1182/blood-2008-03-143354.

28. Mleczko-Sanecka, K.; Casanovas, G.; Ragab, A.; Breitkopf, K.; Muller, A.; Boutros, M.; Dooley, S.; Hentze, M.W.; Muckenthaler, M.U. SMAD7 controls iron metabolism as a potent inhibitor of hepcidin expression. Blood 2010, 115, 2657–2665, doi:10.1182/blood-2009-09-238105.

29. Xu, H.; He, J.H.; Xiao, Z.D.; Zhang, Q.Q.; Chen, Y.Q.; Zhou, H.; Qu, L.H. Liver-enriched transcription factors regulate microRNA-122 that targets CUTL1 during liver development. Hepatology 2010, 52, 1431–1442, doi:10.1002/hep.23818.

30. Li, Z.Y.; Xi, Y.; Zhu, W.N.; Zeng, C.; Zhang, Z.Q.; Guo, Z.C.; Hao, D.L.; Liu, G.; Feng, L.; Chen, H.Z.;, et al. Positive regulation of hepatic miR-122 expression by HNF4alpha. J Hepatol 2011, 55, 602–611, doi:10.1016/j.jhep.2010.12.023.

31. Ovcharenko, I.; Nobrega, M.A.; Loots, G.G.; Stubbs, L. ECR Browser: a tool for visualizing and accessing data from comparisons of multiple vertebrate genomes. Nucleic Acids Res 2004, 32, W280–286, doi:10.1093/nar/gkh355.

32. Fabregat, I.; Caballero-Diaz, D. Transforming Growth Factor-beta-Induced Cell Plasticity in Liver Fibrosis and Hepatocarcinogenesis. Front Oncol 2018, 8, 357, doi:10.3389/fonc.2018.00357.

33. Finnson, K.W.; Almadani, Y.; Philip, A. Non-canonical (non-SMAD2/3) TGF-beta signaling in fibrosis: Mechanisms and targets. Semin Cell Dev Biol 2020, 101, 115–122, doi:10.1016/j.semcdb.2019.11.013.

34. Zhang, K.; Zhang, Q.; Deng, J.; Li, J.; Li, J.; Wen, L.; Ma, J.; Li, C. ALK5 signaling pathway mediates neurogenesis and functional recovery after cerebral ischemia/reperfusion in rats via Gadd45b. Cell Death Dis 2019, 10, 360, doi:10.1038/s41419-019-1596-z.

35. Guidotti, L.G.; Borrow, P.; Brown, A.; McClary, H.; Koch, R.; Chisari, F.V. Noncytopathic clearance of lymphocytic choriomeningitis virus from the hepatocyte. J Exp Med 1999, 189, 1555–1564, doi:10.1084/jem.189.10.1555.

36. Emig, D.; Salomonis, N.; Baumbach, J.; Lengauer, T.; Conklin, B.R.; Albrecht, M. AltAnalyze and DomainGraph: analyzing and visualizing exon expression data. Nucleic Acids Res 2010, 38, W755–762, doi:10.1093/nar/gkq405.

37. Sticht, C.; De La Torre, C.; Parveen, A.; Gretz, N. miRWalk: An online resource for prediction of microRNA binding sites. PLoS One 2018, 13, e0206239, doi:10.1371/journal.pone.0206239.

38. Ge, S.X.; Jung, D.; Yao, R. ShinyGO: a graphical gene-set enrichment tool for animals and plants. Bioinformatics 2020, 36, 2628–2629, doi:10.1093/bioinformatics/btz931.

39. De Smet, V.; Eysackers, N.; Merens, V.; Kazemzadeh Dastjerd, M.; Halder, G.; Verhulst, S.; Mannaerts, I.; van Grunsven, L.A. Initiation of hepatic stellate cell activation extends into chronic liver disease. Cell Death Dis 2021, 12, 1110, doi:10.1038/s41419-021-04377-1.

40. Nakamura, M.; Kanda, T.; Sasaki, R.; Haga, Y.; Jiang, X.; Wu, S.; Nakamoto, S.; Yokosuka, O. MicroRNA-22 Inhibits the Production of Inflammatory Cytokines by Targeting the PKR Activator PACT in Human Hepatic Stellate Cells. PLoS One 2015, 10, e0144295, doi:10.1371/journal.pone.0144295.

41. Colucci, S.; Altamura, S.; Marques, O.; Dropmann, A.; Horvat, N.K.; Mudder, K.; Hammad, S.; Dooley, S.; Muckenthaler, M.U. Liver Sinusoidal Endothelial Cells Suppress Bone Morphogenetic Protein 2 Production in Response to TGFbeta Pathway Activation. Hepatology 2021, 74, 2186–2200, doi:10.1002/hep.31900.

42. Yin, S.; Fan, Y.; Zhang, H.; Zhao, Z.; Hao, Y.; Li, J.; Sun, C.; Yang, J.; Yang, Z.; Yang, X.;, et al. Differential TGFbeta pathway targeting by miR-122 in humans and mice affects liver cancer metastasis. Nat Commun 2016, 7, 11012, doi:10.1038/ncomms11012.

43. Nair, B.; Nath, L.R. Inevitable role of TGF-beta1 in progression of nonalcoholic fatty liver disease. J Recept Signal Transduct Res 2020, 40, 195–200, doi:10.1080/10799893.2020.1726952.

44. Cheung, O.; Puri, P.; Eicken, C.; Contos, M.J.; Mirshahi, F.; Maher, J.W.; Kellum, J.M.; Min, H.; Luketic, V.A.; Sanyal, A.J. Nonalcoholic steatohepatitis is associated with altered hepatic MicroRNA expression. Hepatology 2008, 48, 1810–1820, doi:10.1002/hep.22569.

45. Giannelli, G.; Villa, E.; Lahn, M. Transforming growth factor-beta as a therapeutic target in hepatocellular carcinoma. Cancer Res 2014, 74, 1890–1894, doi:10.1158/0008-5472.CAN-14-0243.

46. Argentou, N.; Germanidis, G.; Hytiroglou, P.; Apostolou, E.; Vassiliadis, T.; Patsiaoura, K.; Sideras, P.; Germenis, A.E.; Speletas, M. TGF-beta signaling is activated in patients with chronic HBV infection and repressed by SMAD7 overexpression after successful antiviral treatment. Inflamm Res 2016, 65, 355–365, doi:10.1007/s00011-016-0921-6.

47. Kordes, C.; Sawitza, I.; Gotze, S.; Herebian, D.; Haussinger, D. Hepatic stellate cells contribute to progenitor cells and liver regeneration. J Clin Invest 2014, 124, 5503–5515, doi:10.1172/JCI74119.

48. Thenappan, A.; Li, Y.; Kitisin, K.; Rashid, A.; Shetty, K.; Johnson, L.; Mishra, L. Role of transforming growth factor beta signaling and expansion of progenitor cells in regenerating liver. Hepatology 2010, 51, 1373–1382, doi:10.1002/hep.23449.

49. Nygard, I.E.; Mortensen, K.E.; Hedegaard, J.; Conley, L.N.; Kalstad, T.; Bendixen, C.; Revhaug, A. The genetic regulation of the terminating phase of liver regeneration. Comp Hepatol 2012, 11, 3, doi:10.1186/1476-5926-11-3.

50. Kojima, S.; Gatfield, D.; Esau, C.C.; Green, C.B. MicroRNA-122 modulates the rhythmic expression profile of the circadian deadenylase Nocturnin in mouse liver. PLoS One 2010, 5, e11264, doi:10.1371/journal.pone.0011264.

51. Gatfield, D.; Le Martelot, G.; Vejnar, C.E.; Gerlach, D.; Schaad, O.; Fleury-Olela, F.; Ruskeepaa, A.L.; Oresic, M.; Esau, C.C.; Zdobnov, E.M.;, et al. Integration of microRNA miR-122 in hepatic circadian gene expression. Genes Dev 2009, 23, 1313–1326, doi:10.1101/gad.1781009.

52. Benes, V.; Collier, P.; Kordes, C.; Stolte, J.; Rausch, T.; Muckentaler, M.U.; Haussinger, D.; Castoldi, M. Identification of cytokine-induced modulation of microRNA expression and secretion as measured by a novel microRNA specific qPCR assay. Sci Rep 2015, 5, 11590, doi:10.1038/srep11590.

53. Hellemans, J.; Mortier, G.; De Paepe, A.; Speleman, F.; Vandesompele, J. qBase relative quantification framework and software for management and automated analysis of real-time quantitative PCR data. Genome Biol 2007, 8, R19, doi:10.1186/gb-2007-8-2-r19.

54. Schlotter, Y.M.; Veenhof, E.Z.; Brinkhof, B.; Rutten, V.P.; Spee, B.; Willemse, T.; Penning, L.C. A GeNorm algorithm-based selection of reference genes for quantitative real-time PCR in skin biopsies of healthy dogs and dogs with atopic dermatitis. Vet Immunol Immunopathol 2009, 129, 115–118, doi:10.1016/j.vetimm.2008.12.004.

55. Livak, K.J.; Schmittgen, T.D. Analysis of relative gene expression data using real-time quantitative PCR and the 2(-Delta Delta C(T)) Method. Methods 2001, 25, 402–408, doi:10.1006/meth.2001.1262.

56. Mestdagh, P.; Van Vlierberghe, P.; De Weer, A.; Muth, D.; Westermann, F.; Speleman, F.; Vandesompele, J. A novel and universal method for microRNA RT-qPCR data normalization. Genome Biol 2009, 10, R64, doi:10.1186/gb-2009-10-6-r64.

57. Klingmuller, U.; Bauer, A.; Bohl, S.; Nickel, P.J.; Breitkopf, K.; Dooley, S.; Zellmer, S.; Kern, C.; Merfort, I.; Sparna, T.;, et al. Primary mouse hepatocytes for systems biology approaches: a standardized in vitro system for modelling of signal transduction pathways. Syst Biol (Stevenage) 2006, 153, 433–447, doi:10.1049/ip-syb:20050067.

58. Boussif, O.; Lezoualc’h, F.; Zanta, M.A.; Mergny, M.D.; Scherman, D.; Demeneix, B.; Behr, J.P. A versatile vector for gene and oligonucleotide transfer into cells in culture and in vivo: polyethylenimine. Proc Natl Acad Sci U S A 1995, 92, 7297–7301, doi:10.1073/pnas.92.16.7297.

59. Nagy, A.; Lanczky, A.; Menyhart, O.; Gyorffy, B. Validation of miRNA prognostic power in hepatocellular carcinoma using expression data of independent datasets. Sci Rep 2018, 8, 9227, doi: 10.1038/s41598-018-27521-y.

