## Supplementary Figures for "Differential modulation of miR-122 transcription by TGFβ1/BMP6: implications for nonresolving inflammation and hepatocarcinogenesis"

Mirco Castoldi

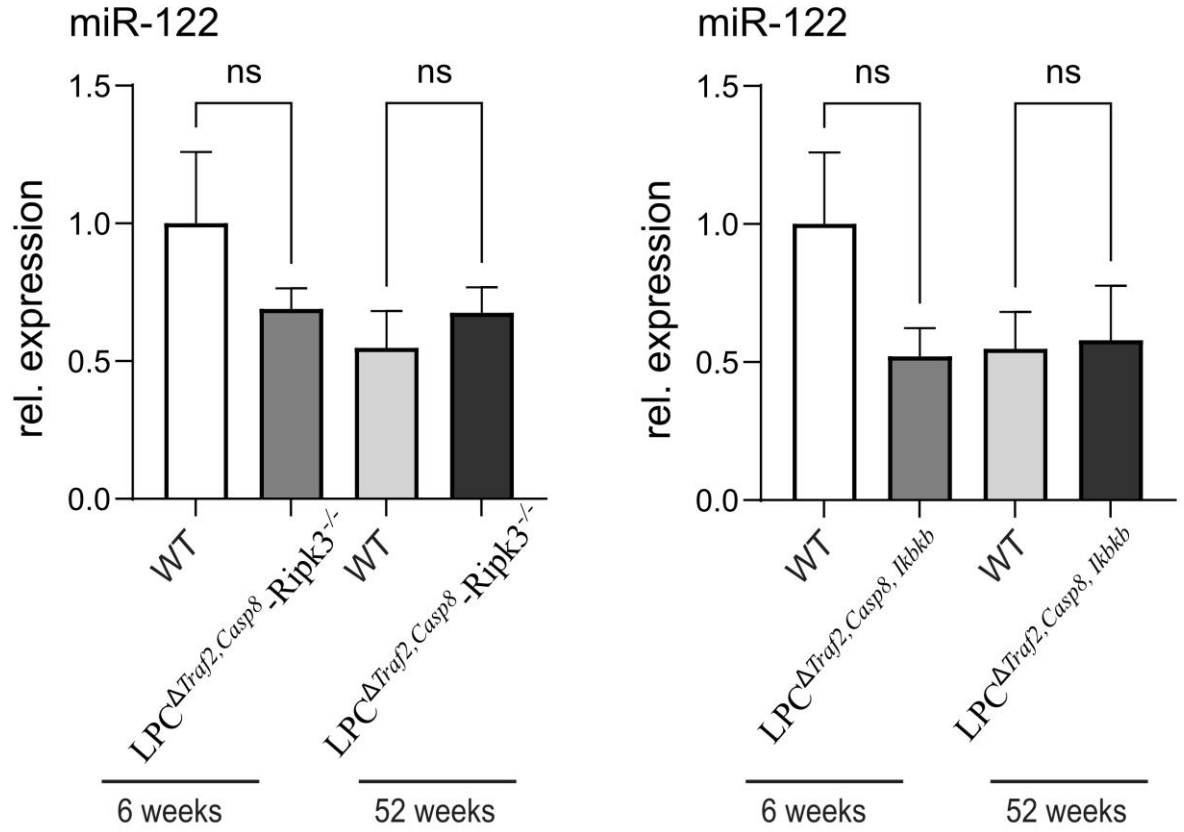


**Supplementary Figure S1: Expression of miR-122 in the livers of LPC$^{\Delta \textit{Traf2, Casp8}}$ –Ripk3$^{-/-}$ and LPC$^{\Delta \textit{Traf2, Casp8,Ikbkb}}$ KO mice**

qPCR analysis of miR-122 expression in the livers of LPC$^{\Delta \textit{Traf2, Casp8}}$ –Ripk3$^{-/-}$ and LPC$^{\Delta \textit{Traf2, Casp8,Ikbkb}}$ transgenic animals (n = 4) and matching controls (n = 4) at 6 and 52 weeks of age. Statistical analysis was carried out by t-test (ns = Not significant).

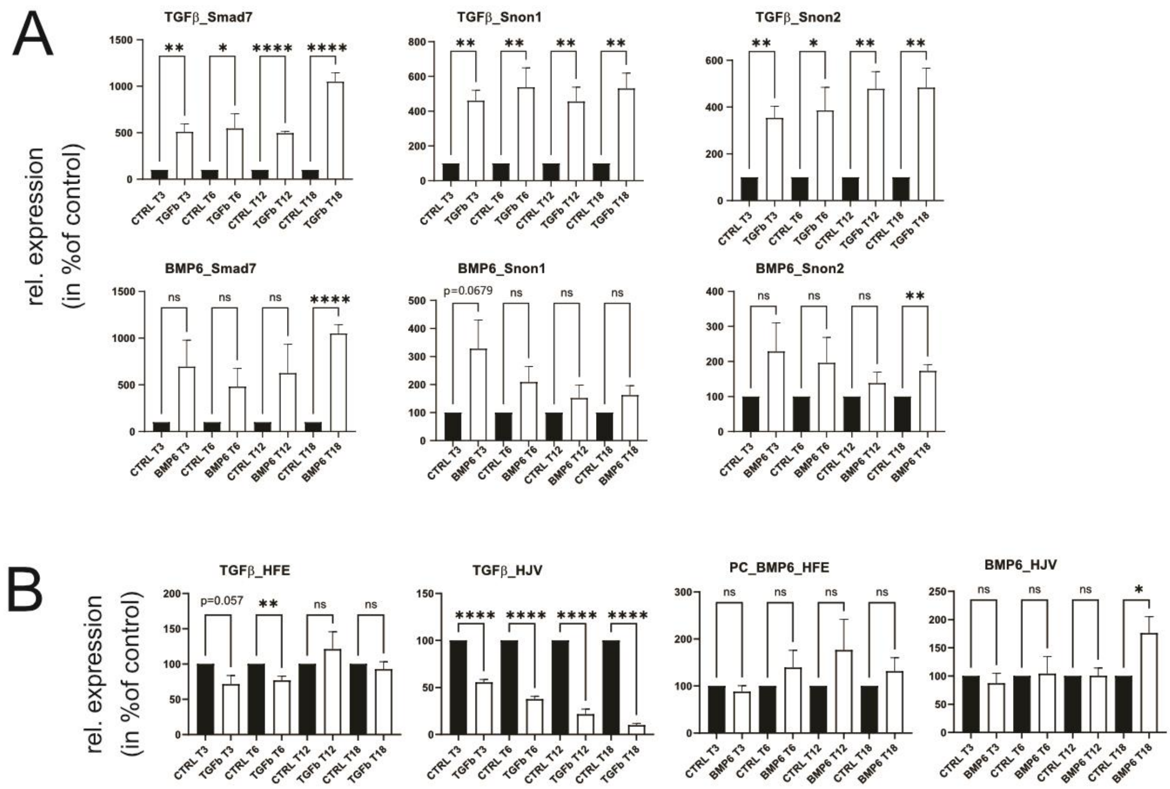


**Supplementary Figure S2: Administration of TGFβ activates the expression of Smad7 and Smad7-responsive genes**

(**A**) qPCR analysis of Smad7, Snon1 and Snon2 expression in primary mouse hepatocytes stimulated with either TGFβ (5 ng/mL, **Top**) or BMP6 (50 ng/mL, **Botton**) (n = 4). (**B**) qPCR analysis of Hfe and Hjv expression in primary mouse hepatocytes stimulated with either TGFβ (5 ng/mL, **Left**) or BMP6 (50 ng/mL, **right**) (n = 4). Data are represented as mean ± SD (n = 4). Statistical analysis was carried out by t-test (*, $p \leq 0.05$; **, $p \leq 0.01$; ***, $p \leq 0.001$; ****, $p \leq 0.0001$).

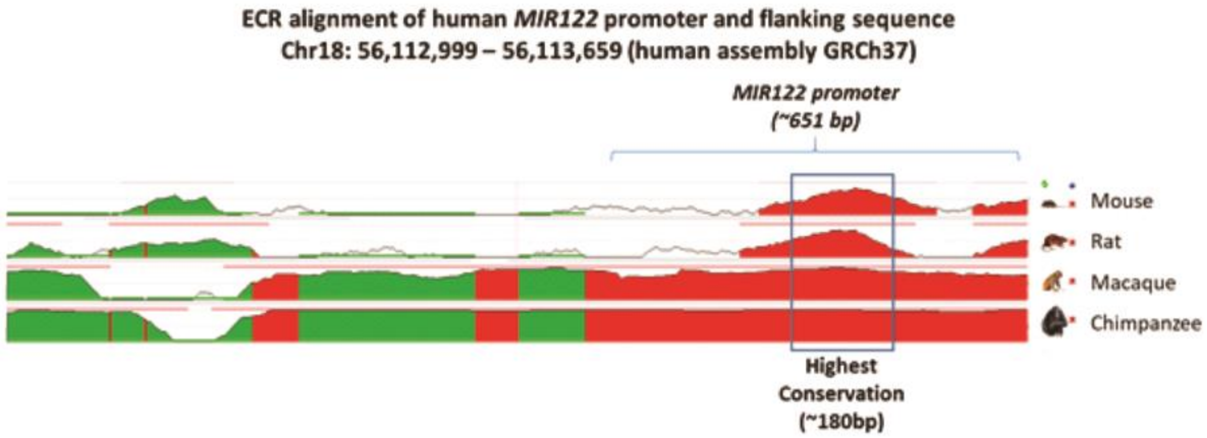


**Supplementary Figure S3: miR-122 promoter, flanking conserved across mammals**

Evolutionary Conserved Regions (ECRs) alignment of the human *MIR122* promoter and flanking regions in mammals.

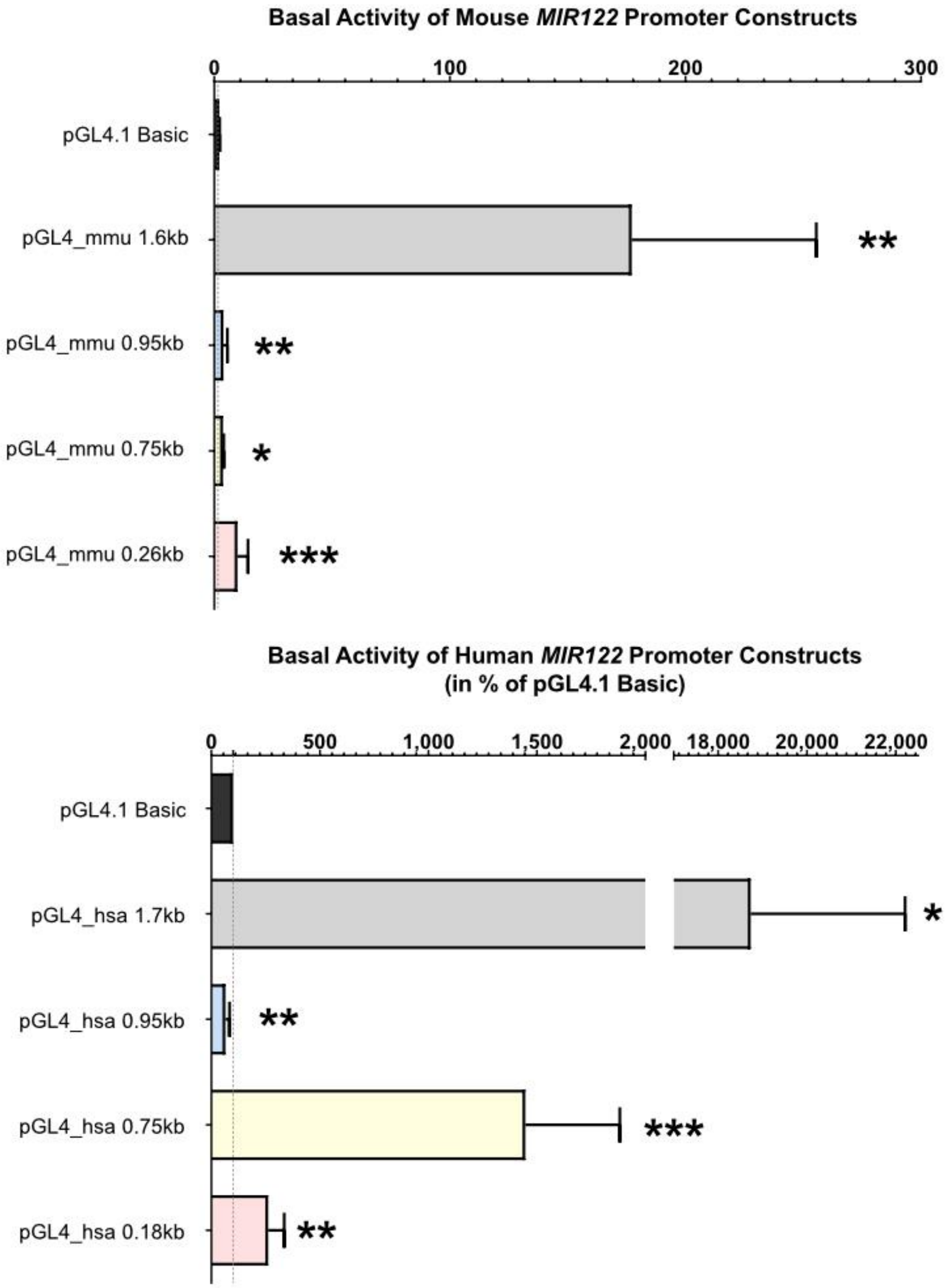


**Supplementary Figure S4: Basal activity of the mouse and human miR-122 promoters**

(**Top panel**) Cells were transiently co-transfected with constructs carry different lengths of the mouse *MIR122* promoter in the pGL4.1-basic vector, containing the promoter-less sequence of firefly luciferase and the pRL vector expressing for the Renilla luciferase as normalizer. After transfection, cells were cultured for 48 h in serum-free medium, and luciferase activity was. The mean ratio of Firefly/Renilla luciferase is presented. Values are illustrated as relative expression in percentage of control ± SD (n = 4). Statistical analysis was performed by two-tailed unpaired student´s t-test (*, $p \leq 0.05$; **, $p \leq 0.01$; ***, $p \leq 0.001$).

(**Lower panel**) Promoter constructs encoding different lengths of the human *MIR122* promoter were generated and cloned into a promoter-less luciferase reporter plasmids (pGL4.1 Basic). Basal promoter activities were measured 48 hours after transfection by quantifying luciferase activity. Constructs containing the shortest MIR122 construct (pGL4_hsa 0.18kb) as well as constructs encompassing the core promoter and genomic flanking sequence (pGL4_hsa 0.75kb and pGL4_hsa 1.7kb) showed an increased basal activity compared to promoter-less basic vector pGL4.1 Basic, whereas promoter activity raised with the length of the flanking sequence. Data represent average luciferase activity in % of pGL4.1 Basic vector ± SD (n = 5). Statistical analysis was carried out by t-test (*, $p \leq 0.05$; **, $p \leq 0.01$; ***, $p \leq 0.001$).

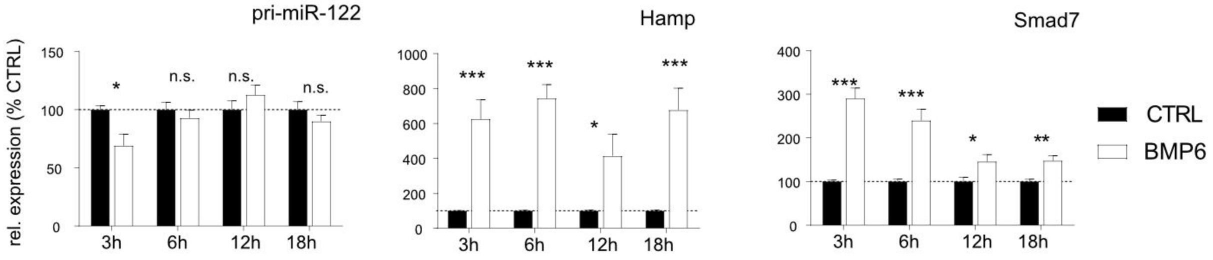


**Supplementary Figure S5: BMP6 administration to Huh-7 cells for 3, 6, 12 and 18 hours**

Effect of BMP6 stimulation on pri-miR-122, Hamp, Smad7 and expression in Huh-7 human-derived hepatoma cells, 3 hrs (n = 8), 6 hrs (n = 8), 12 hrs (n = 6) and 18 hrs (n = 6) after stimulation (Normalized to GAPDH). Results are represented in mean ± SD. Significant differences were evaluated by t-test (*, $p \leq 0.05$).

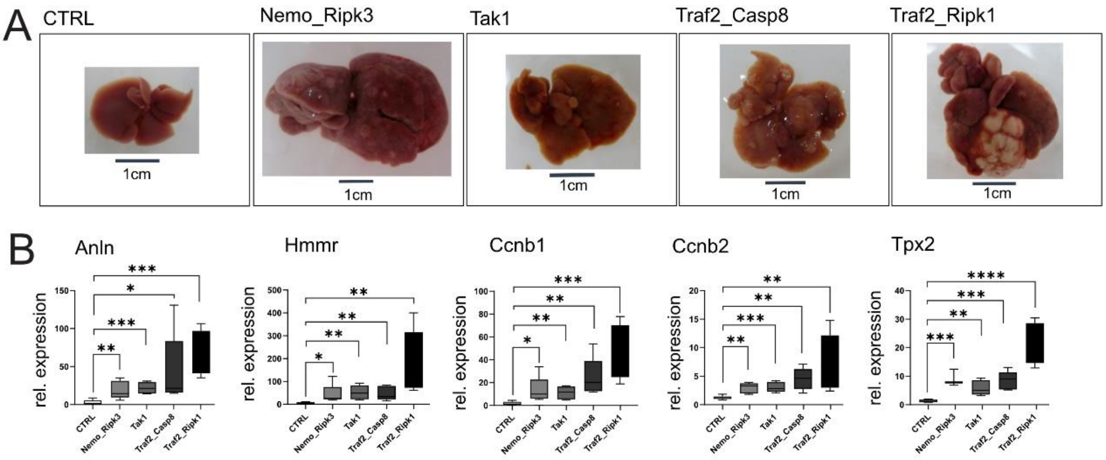


**Supplementary Figure S6 Expression of miR-122 responsive genes in the livers of different mouse model of HCC**

(**A**) Representative pictures of the livers of the different mouse models used in the analysis of miR-122 and of the selected genes. (**B**) qPCR analysis of the expression of Anln, Hmmr, Ccnb1 , Ccnb2, and Tpx2 in the livers of different transgenic animals that autonomously develop liver cancer. Five animals were analyzed for every considered group. Data are represented as mean ± SD. Significant differences were evaluated by t-test by comparing the expression of the GOIs in individual transgenic animal to the CTRL group, *, $p \leq 0.05$; **, $p \leq 0.01$; ***, $p \leq 0.001$; ****, $p \leq 0.0001$).

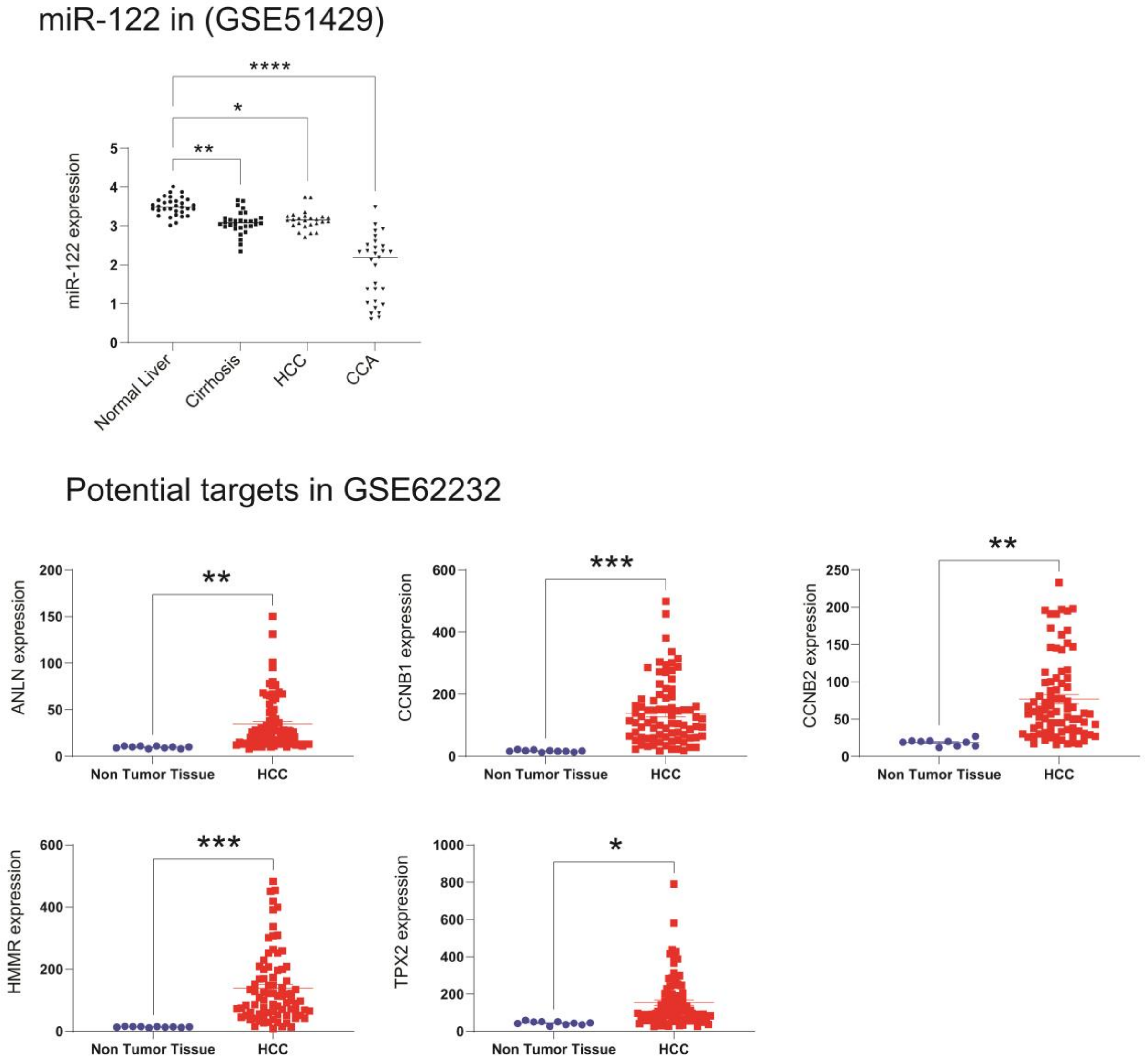


**Supplementary Figure S7: Expression of miR-122 and predicted targets in human HCC samples**

(Top panel) expression of miR-122 in the liver of large cohorts of cancer patients (GSE51429). (Lower panel) Expression of selected genes in the tumor tissue of large cohorts of HCC patients (GSE62232).

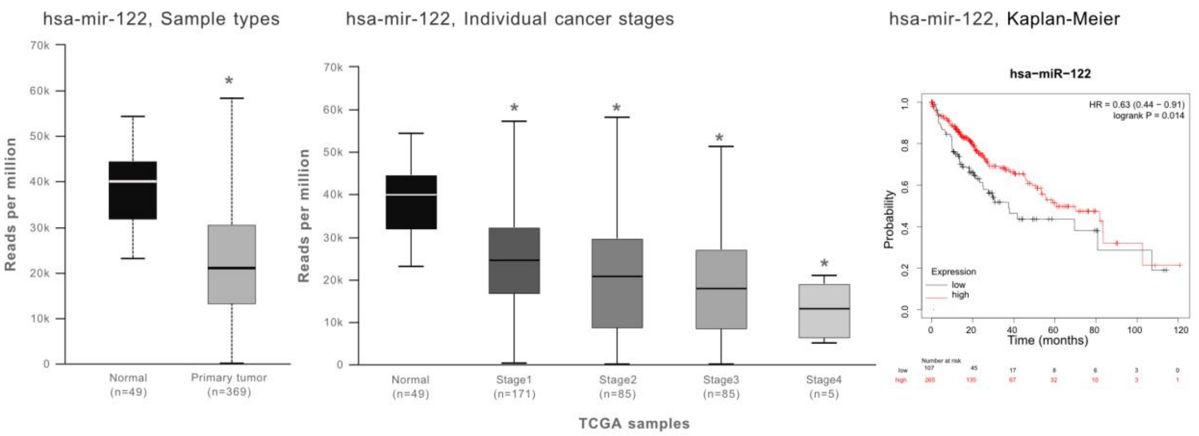


**Supplementary Figure S8: Expression of miR-122 from TCGA samples**

Analysis of miR-122 expression levels in TCGA from cohorts of human patients with LIHC, in total and individual cancer stages as well as Kaplan-Meier from miR-122 in LIHC.

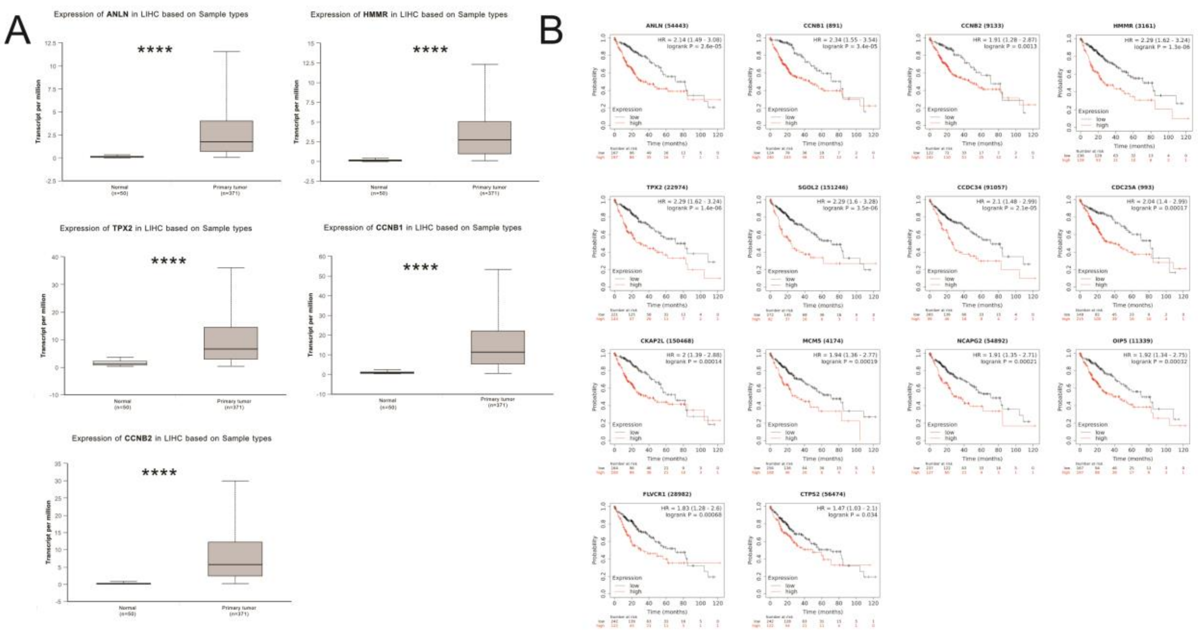


**Supplementary Figure S9: Kaplan-Meier survival curves for selected genes in patients with LIHC**

(A) Analysis of ANLN, HMMR, TPX2, CCNB1, and CCNB2 expression in LIHC cohorts from TCGA data. (B) Analysis of TCGA data from LIHC patients for the 14 predicted miR-122 target genes identified in mouse model of HCC. TCGA data indicates that all the 14 genes that were found significantly upregulated in the mouse HCC model are potential diagnostic marker for human HCC, with lower expression associated with significantly higher survival chance (****, ≤ 0.0001).

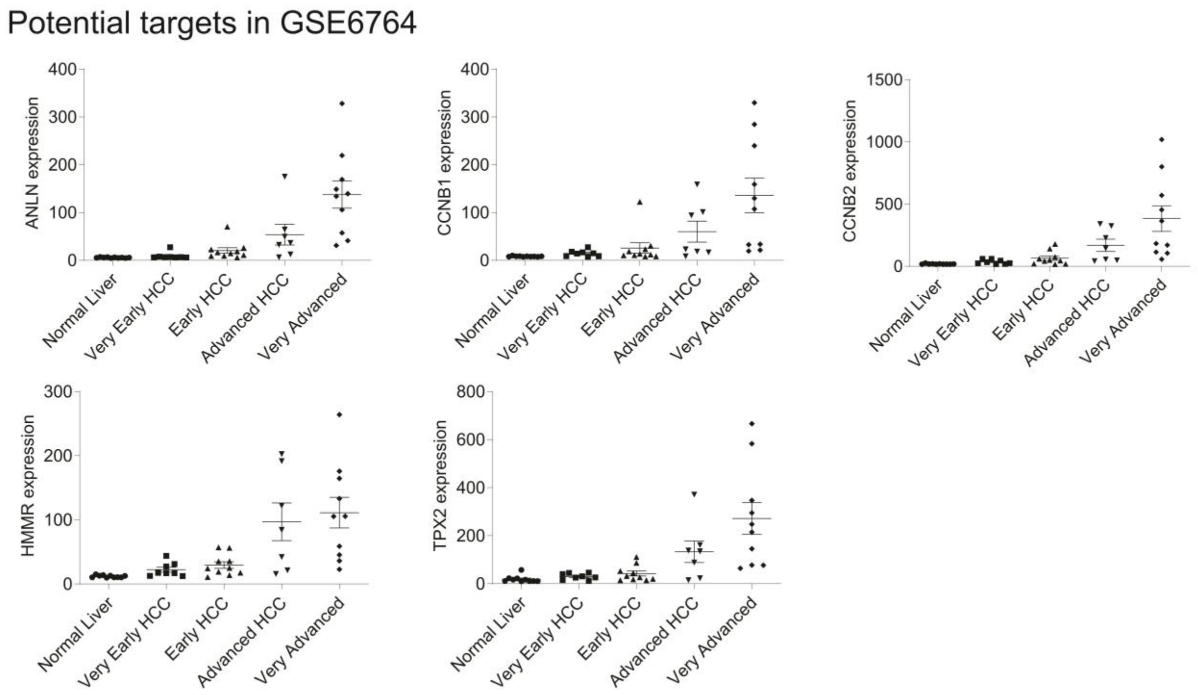


**Supplementary Figure S10: Expression of potential miR-122 targets in large cohorts of human HCC samples**

Expression of selected genes in the tumor tissue of large cohorts of HCC patients subdivided by cancer stages (GSE6764).

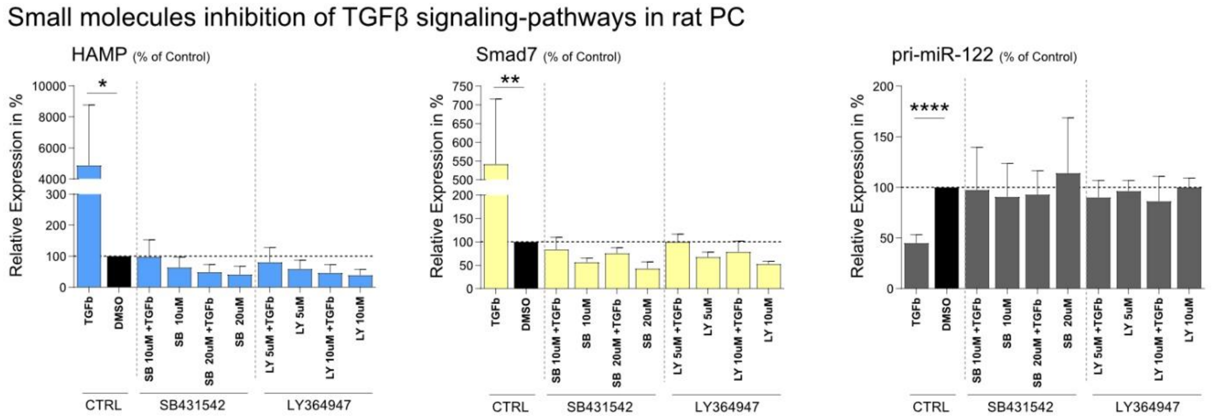


**Supplementary Figure S11: Small molecules inhibition of TGFβ signaling-pathways in rat primary hepatocytes**

Serum starved primary rat hepatocytes were stimulated with 5 ng/ml TGFβ in either presence or absence of the TGFβ inhibitors SB431542 and LY364947 for 3 hours. Control cells were kept under serum-free condition. Following RNA isolation, levels of, Hepcidin (Hamp), Smad7 and primary miR-122 transcript were measured by means of qPCR (Normalized to UbC and Ywhaz). Values are illustrated as relative expression in percentage of control ± SD (n = 3). Statistical analysis was performed by two-tailed unpaired student´s t-test (*, $p \leq 0.05$, **, $p \leq 0.01$).
